# Environmental variability alters and obscures fisheries-induced evolution

**DOI:** 10.64898/2026.10.06.757175

**Authors:** Matthew P. Schumm, Daniel K. Okamoto

## Abstract

Shifts in predation and human harvest often coincide with changes in life history trait distributions in prey populations. These changes may arise via evolution or plasticity, and extrinsic environmental effects may affect such traits. This complexity prompts questions about how trait distributions are shaped by multiple factors’ interactions. We hypothesized that food availability and climate affect both the magnitude, and mechanism (evolution vs. plasticity), by which traits change in response to harvest’s intensity and size selectivity. We tested these hypotheses using individual-based bioenergetic population models allowing both plasticity and evolution. We exposed genetically diverse fish populations to gradients of productivity and stochasticity in food availability, crossed with gradients of selective or nonselective fishing mortality. Food availability shaped traits both plastically and evolutionarily, and modulated how harvesting influenced those responses. Food effects can suppress harvest impacts altogether, or obscure evolution’s effects on phenotypes. However, when climatic factors shape food-driven changes by affecting both fish and their prey, either synergistic or antagonistic interactions can emerge between food variation and harvest in determining phenotype distributions. Our research suggests that understanding consumer-resource dynamics, and climatic processes shaping them, is essential to interpreting how harvest can shape traits relevant to population viability and fisheries sustainability.

## Introduction

Top-down and bottom-up forces can rapidly alter life history trait distributions (Forestier et al. 2020). For instance, predation, food availability, and temperature influence fitness, driving the evolution of reproductive and growth traits (Gordon et al. 2009; Grainger and Levine 2022, Czorlich et al. 2022). These shifts occur through both multi-generational evolution, and phenotypic plasticity, and evolution and plasticity may act in opposition or in concert (Potter et al. 2021). As a result, identifying causes of shifting life history traits in nature remains challenging.

Despite these difficulties, understanding how extrinsic environmental factors and top-down predation jointly shape life history traits is an important aim of basic and applied population ecology. Wild fish populations experiencing human harvest and rapid climate change exemplify the difficulty of disentangling trait changes driven by predation (in this case, harvest) from those caused by other environmental factors (e.g., MacDonald et al. 2018; Hay et al. 2019). Life history changes may represent adaptive responses ultimately improving harvested fish populations’ sustainability (Kindsvater and Palkovacs 2017). However, fitness conferred to individuals by a given life history strategy depends on complex tradeoffs. For example, fast and/or sustained growth can lead to large body sizes, altering ecosystem interactions and increasing potential for resource acquisition, reproduction, defense, or competition (Travis and Lotterhos 2013; Edeline and Loeuille 2021). Yet such benefits may never be realized if they only come at older ages while conferring higher mortality, or come with the cost of delayed reproduction (Audzijonyte and Richards 2018). Moreover, environmental changes can alter these trade-offs. Reductions in food or increases in temperature that affect ectotherms’ metabolic demands can decrease maximum possible growth or reproduction (Uusi-Heikkilä et al. 2011; Audzijonyte et al. 2020). Thus, understanding shifts in life history traits under anthropogenic change requires considering how top-down and bottom-up forces each drive both evolutionary and plastic change.

Plasticity in life history occurs across many taxa in response to both bottom-up and top-down forcing (Stearns and Koella 1986; Gotthard and Nylin 1995; Van Kleunen and Fischer 2003; Than et al. 2020; Potter et al. 2021). Harvest and other predation can drive plasticity both directly and indirectly. Some organisms plastically alter life histories in direct response to predator presence. For example, snails, sea ducks, and oysters all plastically alter life histories or growth in direct response to predator presence or cues (Crowl and Covich 1990; Carroll and Clements 2019; Mohring et al. 2023). Additionally, indirect plastic responses to predation can result from reduced conspecific density, such as declining mangrove density leading to faster seedling-to-juvenile progression (López-Hoffman et al. 2007). Such plastic changes may act in the same direction as selection on genotypes. In this case, plasticity may pre-empt a given evolutionary change, producing that change before it could be produced by selection (Fitzpatrick 2012; Kokko et al. 2017). In contrast, plasticity and evolution may act in different directions (Rochet 1998). In such cases, phenotypic change may or may not manifest even in the presence of evolution at the genetic level (e.g., cryptic evolution; Potter et al. 2021). Therefore, phenotypic data alone are insufficient to assess whether shifts result from evolutionary or plastic change (Pinsky et al. 2021).

Large-scale industrial human harvest dramatically alters survivorship and demography of harvested fish populations. Numerous examples document wild harvest correlating with shifts in life history traits (DeMartini et al. 1993; Koslow et al. 1995; Rochet 1998; Planque et al. 2010; Heino et al. 2015). But it is often unclear if such phenotypic changes in the wild represent evolution, plasticity, or both (Travis and Lotterhos 2013). A commonly cited plastic mechanism is that harvest reduces population density which thereby increases per capita resource availability through reduced competition or higher prey density (Hutchings and Kuparinen 2021). Simple mathematical models predict that increases in per capita resources should result in maturity at younger ages and/or bigger sizes (Day and Rowe 2002). Many empirical examples support these hypotheses (Reznick et al. 1993, Mollet et al. 2007). In contrast, harvest induced evolution is expected to lead to earlier maturation but at smaller sizes. Therefore, observations of harvested fish maturing at both earlier age and smaller size are pointed to as evidence for evolution (e.g., DeCelles and Vidal 2020; Retnoningtyas et al. 2023). But several experimental settings and studies present empirical evidence for plastic reduction in age and length at maturity simultaneously (Alm 1949; Sohn 1977; Uusi-Heikkilä et al. 2011). Similarly, a wild observational study of heavily harvested cod suggested a role for plasticity in maturation changes because of lack of genomic evidence of selective sweeps due to harvest (Pinsky et al. 2021, but see Reid et al. 2023 and Therkildsen et al. 2013).

While changes in per-capita resources can directly affect growth and maturation, they may also modulate how these traits are affected by predation or harvest (Gobin et al. 2018, Thambithurai and Kuparinen 2023). Declines in per capita food can alter energetic trade-offs between investment in growth, reproduction, and acquiring food (Caswell 1982). Importantly, food availability can be shaped by both extrinsic resource productivity but also by conspecific density and competition (e.g., Schrader and Travis 2012). However, many studies on evolution due to harvest assume or keep food per capita and other environmental variables constant–but see Lyberger et al. (2021) who manipulated density independently of harvest selection, Diaz Pauli et al. (2023) who let population density vary with harvest, and Rogers et al. (2023) who directly manipulated food. Low food and resulting low body condition partly explained plastic variation in different age-length combinations at which experimental zebrafish matured (Uusi-Heikkila et al. 2011). Thus, food availability can drive plastic changes which may interact with harvest-induced genetic and phenotypic changes. Individual-based modeling has already shown that fisheries-induced life history evolution may be impeded by slowed growth in low-food environments (Gobin et al. 2018). Additionally, genetic data show that food availability alters fishes’ evolutionary trajectories (Diopere et al. 2018; Czorlich et al. 2022). These findings show that theory on maturation timing considering length and age alone, without considering food, cannot account for how maturation changes. They raise largely unanswered questions about common patterns in eco-evolutionary responses to co-occurring harvest and food fluctuations.

Apart from food availability, many other extrinsic conditions may affect fish life history trait distributions, and these may simultaneously affect all species in the community (Piou and Prévost 2013; Hiltunen et al. 2018). Changes in temperature are expected to affect species across trophic levels. In such cases, impacts of harvest on individual traits have the potential to be modulated simultaneously by temperature effects on multiple species as they interact (Dee et al. 2020). Moreover, a given temperature change may similarly affect predators and prey or it may oppositely affect them (e.g., positively for prey, but negatively for predators). This complexity means that understanding how temperature variation shapes trait distributions requires understanding both direct physiological effects of temperature on a focal population and the indirect effects of temperature on its food sources (Lindmark et al. 2022).

In our study, we address how shifts in mortality and variation in both food levels and temperature interactively affect a population’s trait distributions both evolutionarily and plastically. Using individual-based bioenergetic models of population dynamics (Figure 1), we manipulated harvest rate and selectivity, base productivity level and its variability, and temperature effects on fish and their food. To separate evolutionary from plastic contributions, we simulated scenarios with and without heritable trait variation (see Table 1 for model characteristics and environmental variables manipulated). We first tested whether harvest of a fish population, and trends and variability in their prey, favor evolution of faster life history (Dunlop et al. 2009). Additionally, we tested how results change when these dynamics are affected by temperature that can similarly or oppositely affect fish and their prey. We compared results using models of a herring (Clupeidae)-like fish and a cod (Gadidae)-like fish. Compared to gadids, clupeids generally are much smaller, mature earlier, and have higher adult mortality. These differences affect the selection they experience under changing mortality regimes (Kindsvater et al. 2016; Schmid et al. 2022). First, we hypothesized that fluctuations in food reduce the fitness benefits of an accelerated life history in response to harvest, and therefore reduce harvest-induced evolution. Such reduction could be caused by interactions between harvest’s selective effects (if any), and food availability conditions (Gobin et al. 2018; Thambithurai and Kuparinen 2023). Second, we hypothesized that fluctuations in food reduce overall phenotypic change in life history due to harvest because of their strong plastic effects on life history (Audzijonyte and Richards 2018). We also explored how trait distributions change with simultaneous temperature effects on both a fish and its food source, as contrasted with varying food directly. Supplemental Table 1 includes more information about hypothesized effects. Our approach allows us to identify, across broad life history strategies, how and when harvest-induced change in trait distributions can occur because of plasticity or evolution, and how such change is affected by food stochasticity and other possible climate effects.

**Figure 1.**
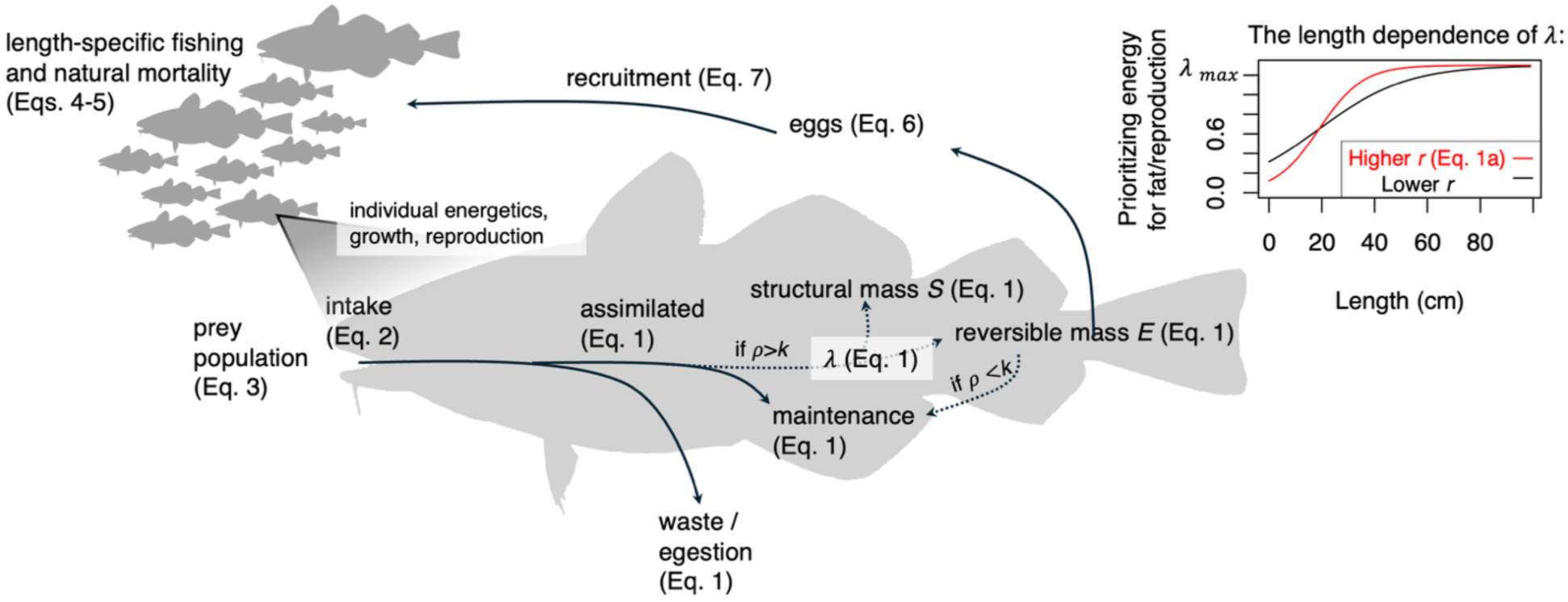
The overall structure of and processes represented in the model. These include (1) the individual energetics, growth, and reproduction of each individual fish, (2) the dynamics of the population through recruitment and size-specific natural mortality and recruitment with heritable values of *r*, and (3) the dynamics of the prey population. Dotted lines show energy flows that only occur under certain conditions, depending on the condition of the fish. The “length dependence of *λ*” inset shows the resource allocation curve (Equation 1a) which is the trait that is allowed to vary genetically between individuals in the model. There is a “small-switching” life history curve in red to show the energy allocation pattern with length of a fish with a high value of *r*, and a curve in black to show the energy allocation pattern with length of a fish with a lower value of *r*.

**Table 1.** Focal model parameters that were varied for testing the hypotheses described in Supplemental Table 1. These are the parameters which are described as having been varied for constructing the different experimental crosses described in the Methods section: “Simulation experiment design”.

| Parameter | Parameter description | Levels | Description of levels |
| --- | --- | --- | --- |
| $F_{max}$ | **Fishing mortality**<br>**(maximum)** | 14 levels (0 to 1.8) | Low (none) to high fishing mortality.<br>Fully crossed with $CV_R$ in simulations carried out, with other parameters at reference values. |
| $z_f$ | **Fishing selectivity** | 2 levels (0.005, 0.5),<br>reference value 0.5<br>(selective harvest) | The probability of harvest changes <5%<br>over the fish’s potential range in length<br>over lifetime, vs. probability of harvest<br>goes from ~0% at initial length to<br>maximum fishing mortality at largest<br>possible length. Crossed with all levels<br>of $F_{max}$ in Figure 4 with other params.<br>held constant at reference level. |
| $CV_R$ | **Food variability** | 14 levels (0 to 8),<br>12 levels of 0 to 2<br>for noise in mean<br>temperature | Equal to $\sqrt{e^{\sigma_R^2} - 1}$ where $\sigma_R$ is the<br>standard deviation for a Normal<br>distribution used to generate noise,<br>which is then colored and scaled to a<br>lognormal distribution. Fully crossed<br>with $F_{max}$ in simulations carried out,<br>with other parameters at reference<br>values. |
| $K_R$ | **Base food productivity level** | 2 levels (3e11 or 6e10 for cod, 3e14 or 6e13 for herring); higher values are the reference levels | Two different orders of magnitude of carrying capacity of the food resource in the absence of its predator (the fish), a low one at which fish are affected in growth by population density, and a high value at which they are not. Crossed with all levels of $F_{max}$ in Figure 4 with other params. held constant at reference level. |
| $V_G$ | **Starting genetic variance in trait** | 2 levels (population variance of 0 in uniform distribution of initial assigned genotype values, variance of ~12 with a mean of 6). Reference level of 12. | No genetic variance (and thus no heritability), vs. some. Fully crossed with $CV_R$ and $F_{max}$ in simulations carried out, with other parameters at reference values. |

## Methods

### Model overview

To assess how length-selective harvest and stochasticity in food availability affect life history via evolution and plasticity, we developed an individual-based, size structured model. We allowed traits such as length at maturity and maximum size to vary as emergent properties, by directly modeling underlying bioenergetic processes, rather than making those traits directly heritable. To simulate dynamics of populations, we used a superindividual-based model (*sensu* Eikeset et al. 2016). Instead of tracking individuals separately, the model tracked groups of fish sharing a cohort and phenotype (’cohort-phenotype combinations’ or all fish with a particular value of the trait *r* - see the Figure 1 inset, and Figure 2).

**Figure 2.**
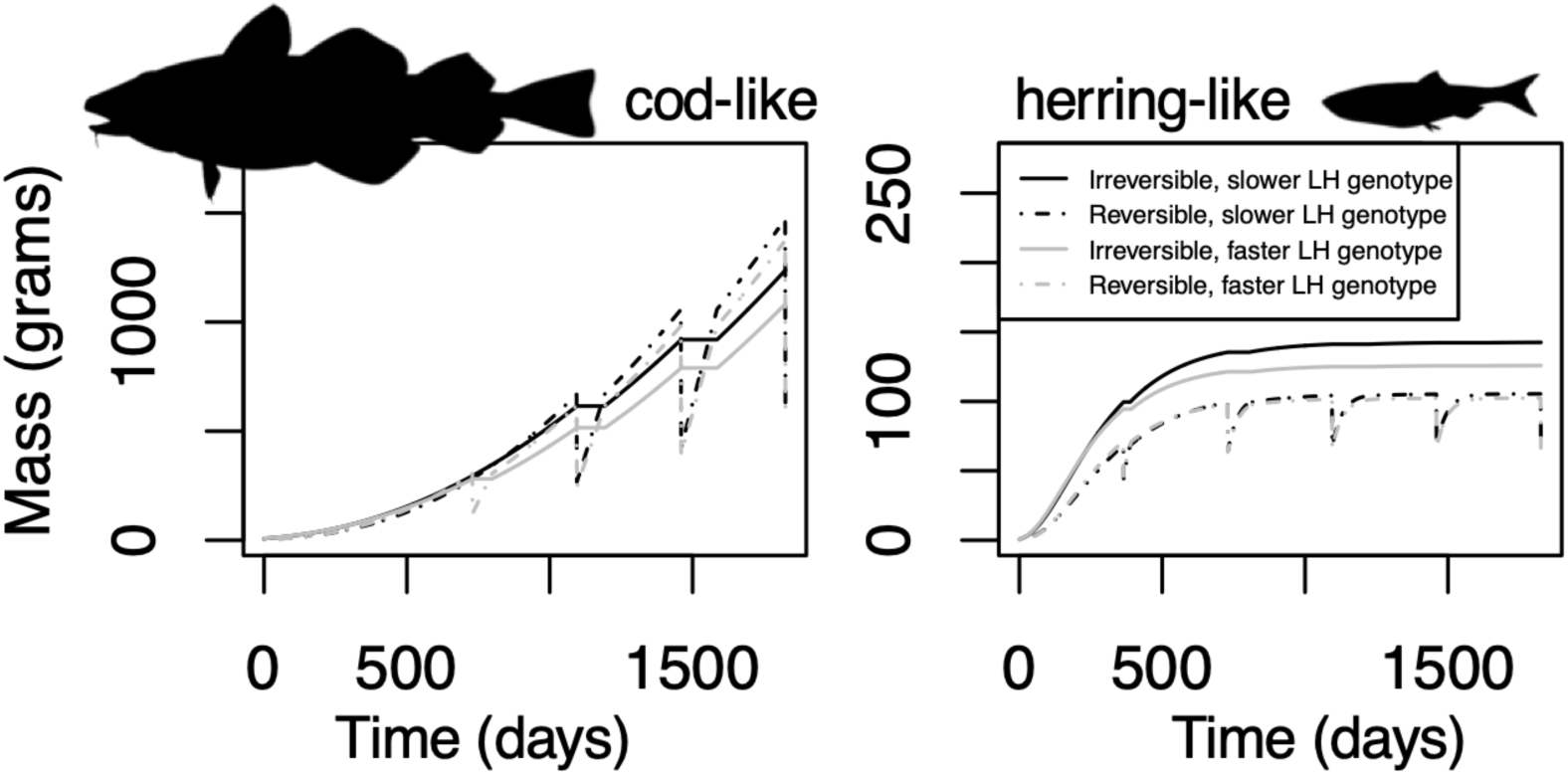
These figures show the differences between fish in the model with different genetically-determined bioenergetic phenotypes. They show the growth of reversible and structural mass in the two modeled fish species, given unlimited food resources, for fast- and slow-life-history genotypes (in the first five years of life). The solid lines show the growth over time since hatching in irreversible, structural mass for fish with different genotypically influenced values of the parameter *r* (Supplemental Table 2). The dotted lines show the growth over time in reversible mass for fish with those two different *r* values. The effects of these different *r* values on resource allocation are those shown in Figure 1, in the “length dependence of *λ*” inset. Grey lines are the growth in these two biomass pools for the high-*r*, “small-switching” fish, and black lines are the growth for the low-*r* fish. The grey lines can be seen to be affected more and earlier by reproduction than the black.

The key evolvable life-history trait in our model was the rate at which fish alter their proportional energy investment into reproduction versus structural growth as they grow larger (represented as *r -* see description below and Supplemental Table 2). A higher value of this parameter *r* leads to earlier and higher fecundity at earlier ages, but also reduces other key traits such as maximum adult length (Figure 2), maximum fecundity, maximum food acquisition, and survival from (non-human) predation. See Audzijonyte and Richards 2018 for more about this parameter. We describe how heritability and variability in *r* is simulated in *Simulating slow and fast life-history genotypes* below.

Our model allowed for plastic variation in life history traits at a given body length by accommodating distinct pools of structural (*S*) and reversible (*E*) body mass consistent with the principles of dynamic energy budget theory (Kooijman 2000) and thus allowing nutritional memory. The flow of energy was modeled from fish consumption and assimilation, to allocation to maintenance, and then to growth of structural and reversible mass. The parameter *r* controlled the rate at which fish shifted from prioritizing structural mass to reversible mass and reproduction as they grew larger. The growth of these pools of body biomass (*E* and *S*) as a function of body size is shown with different values of the *r* parameter and for both cod and herring in Figure 2. This pattern results from the underlying differences in length-specific allocation in Figure 1’s inset plot. Periodic declines in reversible mass *E* (Figure 2) represent the depletion of *E* at each breeding season (e.g., Audzijonyte and Richards 2018). Figure 2 shows hypothetical outcomes when food was unrestricted and under fixed environmental conditions.

However, these trajectories plastically change when food becomes limiting or under different temperature regimes, despite a fixed value of *r*, due to variation in energy available to fish for growth. For example, when food availability becomes limiting, fish are forced to use reversible mass to pay maintenance costs rather than invest in growth of *E* or *S*.

### Individual energetics and growth dynamics

Following dynamic energy budget (DEB) theory (Kooijman 2000), we assume growth rate, as well as fecundity and maturity schedules, are emergent properties of organismal energetics and traits rather than direct phenomenological functions of size or condition.

Prioritization of growth vs. reproduction is determined by traits including *r* (as described above), quantity of energetic reserves, and energetic constants (*λ_opt_* and *λ_max_*). We assumed that the optimal ratio of reversible or reserve mass (*E*) to structural mass (*S*) (*λ_opt_*) increased as animals grew (Jørgensen and Fiksen 2006) up to a fixed maximum value *λ_max_* at large lengths. The rate at which *λ_opt_* approaches *λ_max_* as fish grew was controlled by the rate parameter *r*, and the scale parameter *l̄*. *λ_max_* and *l̄* are both fixed for all fish within a species. This function is shown in Equation 1a:

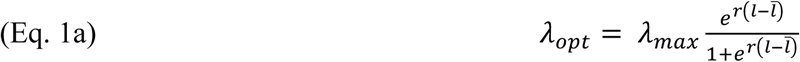

where *r* is a genetically controlled heritable trait parameter.

Animals were able to grow when they had sufficient energy available, where *ρ_net_* was the net energy acquisition after maintenance costs are paid. Fish allocated this energy between *E* and *S* to achieve their length-specific *λ_opt_* ratio. They did so by first allocating energy to *E* until *E*:*S* was equal to *λ_opt_*. The amount by which *E* fell short of the value needed for this length-specific optimal ratio was calculated for the *i*th group of individual fish at time *t* as the following:

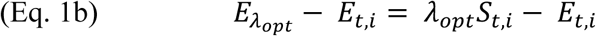

If and when *λ_opt_* was achieved, fish allocated energy between *E* and *S* to maintain their length-fspecific *λ_opt_* ratio according to the following equations:

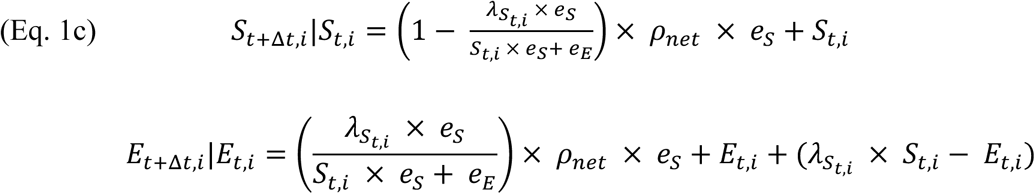

where, and *e_S_* and *e_E_* are the efficiencies with which *ρ_net_* is converted to *S* and *E*, respectively. Note that length varied with structural mass according to a fixed power-law relationship shown in Eqn. 1d:

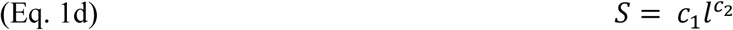

When animals were in energetic deficit (i.e., *ρ_net_* ≤ 0), they drew on their reserves (*E*) to pay costs without any change in *S*. Only when maintenance costs were met was remaining energy intake spent on building tissue.

Before energy could be used for growth, animals needed to first pay energetic maintenance costs associated with existing tissues (i.e. due to activity, repair, etc.). Mass-specific maintenance costs for structural (*S*) and reversible mass (*E*) were defined by *c_S_* and *c_E_*, respectively. These were chosen from the cod model by Audzijonyte and Richards (2018) and from the herring model by Huss et al. (2012). Total maintenance costs *k*, and remaining energy *ρ_net_* following maintenance, are given by the following rates:

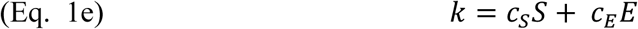

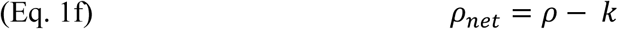

These equations assumed metabolic maintenance and activity costs for the two tissues each grow isometrically. However, for the cod-like model, total body maintenance cost *k* had an emergent allometric scaling exponent of less than 1 because the ratio of reversible mass to structural mass increased as fish grew, and reversible mass has lower mass-specific maintenance costs than structural mass.

An emergent property from our model is that reproduction increases and growth in length slows once a threshold body length has been reached, as observed empirically in fish resource allocation shifts (Policansky 1982; Engelhard and Heino 2004; Arula et al. 2017; Sewall et al. 2019). Slowing of growth in length occurred in our models because (1) fish increasingly prioritize investing in *E* over *S* (higher *λ_opt_* ) as they get longer and (2) maintenance costs increase faster with body size than energy acquisition (higher allometric exponent value for maintenance than food acquisition). Thus, emergent properties from the physiologically structured model were that length-at-age approximated a von Bertalanffy shape, and fecundity at length was hyper-allometric. The change in *λ_opt_* with a fish’s increasing length (Equation 1a) is shown in Figure 1 while the growth of those mass pools given different values of the parameter *r* of this function is shown in Figure 2. These figures illustrate how individuals’ high values for *r* divert resources to energy stores and reproduction at relatively smaller lengths.

### Energy acquisition and prey dynamics

We simulated fish consumption of available prey through a system of ODEs (ordinary differential equations) during a single bout per day. We assumed no growth, birth or mortality of fish during that bout. Consumption rates by each fish cohort (*ρ*) on a given day varied by its structural mass *S*, and by the density of the prey resource (*N*), according to a Holling type II functional response. The prey resource (*N)* also could change during the simulated bout, and it dynamically responded to consumption from each individual fish in the fish population (i.e., all individual fish within each cohort) while growing according to logistic growth:

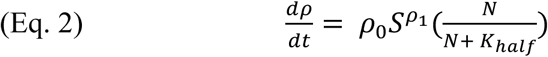

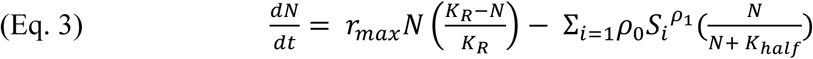

where *K_R_* represents the carrying capacity and *r_max_* denotes the maximum potential growth rate of the prey population, and index *i* represents each individual fish (Royama 2021). *ρ*_0_ and *ρ*_1_are allometric scaling parameters and *K*_*half*_ is the prey density at which consumption is 50% of the asymptotic maximum. We set *ρ*_1_to 2/3, consistent with surface area functions for energy acquisition in energy budget models generally (Von Bertalanffy 1957; Nisbet et al. 2000; Audzijonyte and Richards 2018). Stochastic change in food availability was introduced to the model by allowing the carrying capacity *K_R_* to vary among days via lognormal first order autoregressive process (controlled by the autocorrelation parameter *φ* and coefficient of variation *CV_R_*, Table 1, Supplemental Methods). To simulate reduced base food levels without stochasticity, we set *CV_R_*=0 and compared outcomes when carrying capacity *K_R_* is unchanged vs. reduced by 80%.

### Mortality

Mortality was compounded daily to project the next day’s abundance (*N_l_*_,*E*,*t*+1_ ) of a given cohort-phenotype combination with a given length *l* and reversible mass *E*, at a given initial fish abundance (*N_l_*_,*E*,*t*_ ) and length and condition at the end of the day (*t*):

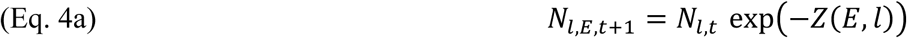

Total mortality was the sum of natural mortality [*M*.(*E*, *l*)] and a length-specific fishing mortality

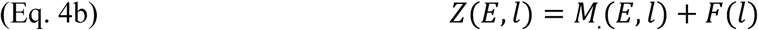

Natural mortality was the sum of a baseline mortality hazard *m*_0_, plus a predation hazard that declined exponentially with length from a maximum *m_p_*, and a condition hazard that declined exponentially with the ratio *E/S,* with predation- and condition-related mortality declining with rates *z_p_* and *z_c_*, respectively.

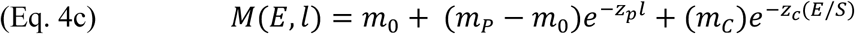

Instantaneous length specific fishing mortality (*F*(*l*)) increased logistically with length:

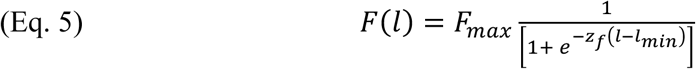

where *l_min_* is the minimum catch size, *z_f_* is the rate and *F_max_* is the instantaneous fishing mortality for fully susceptible fish. Smaller fish are less susceptible to the fishery and thus fishing mortality declines from *F_max_* with smaller fish size via a logistic function with rate *z_f_*, and approaches zero below *l_min_*.

### Fecundity and recruitment

Annual recruitment was modeled as a Ricker density-dependent function determined by female fecundity (assuming a 50:50 sex ratio) and halved to only explicitly count and continue simulating recruits that were female (assuming males and females are equivalent in all respects other than reproduction):

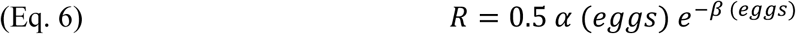

Females reproduced once at the end of the year, representing a single “breeding season” timepoint during the year. Fecundity was proportional to reversible mass (*E*) times the fraction allocated to reproduction, minus non-gametic reproductive overhead (e.g., courtship, migration):

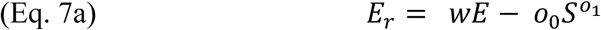

where *o*_0_ and *o*_1_ are allometric parameters for how non-gametic reproductive overhead scales with structural mass (*S*).

Length at maturity in our paper refers to the body length at which a fish was first able to reproduce in its lifetime. Length at maturity, then, is an emergent property from Equation 7a.

Specifically, length at maturity for a given *r* value in a given breeding season is the length of the youngest cohort of fish that can attain *E_R_* > 0. This metric was then summarized across the population in a given year by taking the average of length at maturity across *r* phenotype values, weighting phenotypes by their proportion of the population.

In future years beyond first reproduction, a fish was still only able to reproduce if *E_r_* was positive, otherwise they skipped spawning. Note that skipped spawning was rare for larger animals when food is abundant because the costs scaling exponent *o_1_* (Equation 7a) was <1 and thus costs per unit mass declined with body length (Engelhard and Heino 2005). Skipped spawning could result from low allocation to reversible mass (i.e., the fish is immature) or from having extremely depleted reversible mass due to food shortage.

Egg production is thus:

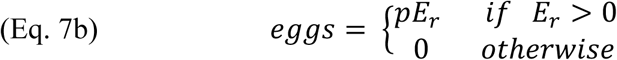

where *p* is the cost of egg production per gram of reserves used.

### Simulating slow and fast life-history genotypes

Genetic variation between individuals was modeled as distinct allele frequencies for each group of fish in each cohort sharing a phenotype. We simulated additive polygenic phenotypes from each of 20 biallelic loci (a typical number of loci for these models, Kuparinen 2021) with no epistasis or linkage. At each locus, there were two alleles (1 or 0). Summing across all 20 loci gives a value for a given haplotype ranging across integers 0 to 20. During reproduction, offspring genotypes were simulated based on the allele frequencies of all mature cohort-phenotype groups contributing gametes. Once offspring were produced and their genotype was determined, we simulated developmental stochasticity and assigned phenotypes based on a truncated Normal distribution (R package ‘truncnorm’, Mersmann et al. 2018) with a mean equal to the average phenotypic value expected for their summed genotype (breeding value, i.e., the aforementioned sum ranging from 0 to 20) and bounds from 0 to 20, and thereafter rounded to the nearest integer. The standard deviation, *σ_devo_* (Table 1), of that distribution for the noise in what actual phenotype for *r* was produced from a given genotype was equal to 1 for most model runs, but was varied in some model runs to assess the sensitivity of model results to amount of heritability in *r*. Heritability was then an emergent function of both (i) genetic diversity, and (ii) any developmental stochasticity, or within-environment variation driving plasticity, that produces distinct phenotypes from the same genotype (Vogt 2015). All individuals sharing a value of *r* within a cohort were grouped and tracked as a “cohort-phenotype combination”, and the allele frequencies of each of these groups of individuals were recorded.

### Temperature-dependency

We also evaluated how extrinsic factors such as temperature may shape plastic and evolutionary changes in traits by affecting both predator and prey. To do so, we modeled positive, monotonic, exponential temperature dependence in the fish’s attack rate (intake parameter *ρ*_0_) and fish maintenance costs (*c_S_* and *c_E_* parameters), as well as its effect on the carrying capacity of the food source *K_R_* (Equation 3) or population growth rate of the food source (*r_max_*). In this way, we model both direct physiological effects of temperature on the fish, and indirect effects of temperature on fish through its prey. These modeling choices were made based on work by Lindmark et al. (2022), a size spectrum model representing a whole food web of organisms in the Baltic Sea, with the fish taxa including cod and herring species. They also modeled temperature dependency of the productivity at the base of the food web with this positive relationship between producers’ growth rate and increasing temperature, and this negative relationship between producers’ carrying capacity and increasing temperature. The parameters we set as temperature-dependent were multiplied by a monotonic Arrhenius function to represent simple temperature dependency of a single reaction rate (assuming we are below any critical thermal limits where the reaction would stop if it were too hot):

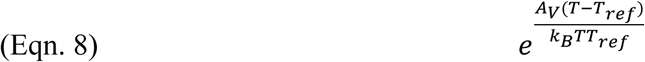

where *A* controls the amount and direction of the parameter’s relationship to temperature, and where this *A* parameter is different for each parameter being multiplied to add temperature dependency. *T_ref_* is a baseline reference temperature, and *k_B_* is Boltzmann’s constant. We simulated temperatures using a log-normal distribution as described above with mean *T_ref_.* We simulated one set of model runs with variation in temperature with a negative relationship between increasing temperature and food source carrying capacity *K_R_*, and another with a positive relationship between food source maximum potential growth rate *r_max_*.

### Simulation experimental design

We used the model to quantify effects of gradients in six key properties (Table 1) for both cod and herring-like models. These included maximum fishing mortality (*F_max_*), food variability (carrying capacity *CV_R_*), base food productivity (*K_R_*), length-selectivity of harvest (*z_f_),* and heritability in energetic allocation (*V_G_*) with details shown in Table 1. For each parameterization we carried out 500 model simulations for 40-year horizons following an introduction of harvest after an initial equilibration of 60 years. We ran a fully crossed set of runs for each combination of *CV_R_*, *F_max_*, and *V_G_* with 500 replicates for each combination of values of those parameters while holding other parameters constant. Table 1 details which other treatment combinations were simulated for key parameters. The initial mean value of the evolvable parameter *r* when evolution was allowed to occur was halfway between the minimum and maximum value in the population, i.e., 0.5 on the scales shown in Figures 3-5. We also conducted sensitivity analyses for parameters including maximum condition-dependent mortality, size-dependent predation mortality, and level of correlation in resource noise, to determine the importance of these factors in model outcomes. Additionally, we ran some versions of the model with genetic diversity in traits other than *r* (but holding *r* constant) including genetic diversity in length-specific ability to acquire food (*ρ*_0_), and genetic diversity in the amount of energy per unit biomass that must be spent on maintenance (*c_S_* and *c_E_*) that can shape genetic differences in growth rate in fish (Uusi-Heikkilä et al. 2011; Auer et al. 2018; Therkildsen et al. 2019). We chose a 40-year horizon as any longer timescales are likely to experience long-period trends, other shocks, and ecological interactions not included in our model. Because evolution had not always reached an equilibrium by the end of this time, we also evaluated some cases for longer or shorter periods to assess any dependence on time horizon of our main results. When modeling temperature effects, we simulated at least 700 model runs for each combination of temperature stochasticity and maximum fishing mortality due to increased outcome variation.

**Figure 3.**
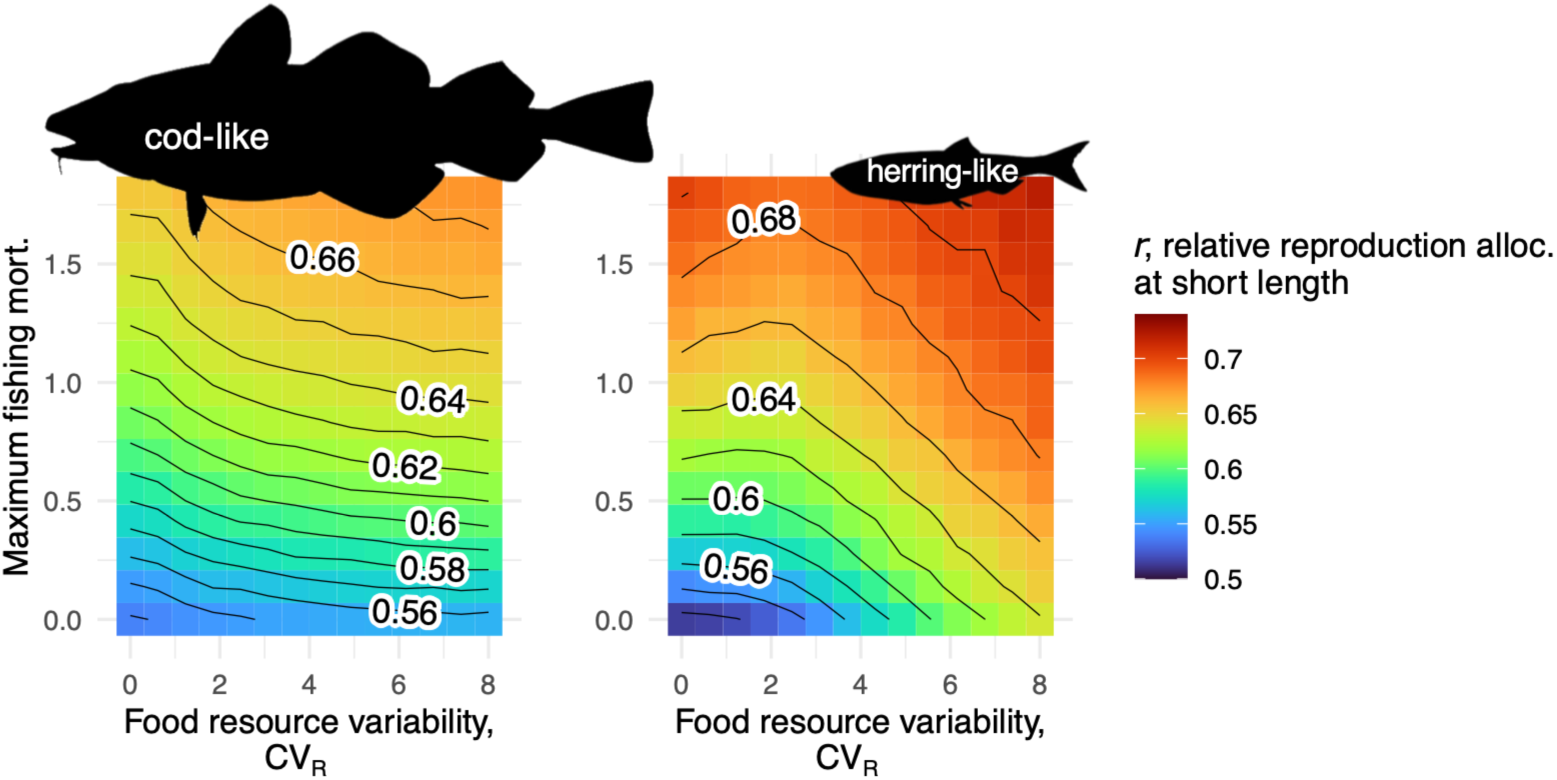
Average evolutionary change of simulated populations under different levels of harvest and food variability after 40 yrs. of harvest. Contours and the colors of the pixels represent the value of *r* (Equation 1a, Supplemental Table 2) on standardized scale where 1 is the maximum value possible and 0 the minimum averaged across all individuals and across 500 model runs each. The x axis shows the size of the coefficient of variation in the temporally autocorrelated random values of the food resource carrying capacity, so larger x values mean more variability imposed in food resource carrying capacity. The y axis shows the value of maximum fishing mortality, F_max_. F_max_ is the maximum possible value of fishing mortality of susceptible fish, where susceptibility is controlled by a length-specific selectivity ogive described in Equation 5. The value of *r* is determined only by genotype and by random developmental stochasticity, so change in the average value of *r* in this model implies genotypic evolution. The starting average value of *r* in the model is halfway between the possible high and low extremes (0.5 on the normalized scale for *r* in Figures 3 and 4, also see *r* in Supplemental Table 2), so values above 0.5 represent an evolutionary increase in *r*. Results are shown for both the cod-like and herring-like models.

We built our model in R (R Core Team 2021), using the deSolve package for differential equation integration. A shinyapps.io link (Technical Supplement I) to code, a detailed description of how the model was coded in R, an interactive model demo, and a Zenodo link to code and simulated data from the model can be found in supplemental material.

## Results

### Life history evolution in response to harvest and food variation

Preferentially harvesting larger fish caused substantial evolutionary increases in length-specific energy allocation to reproduction. However, this evolution was sensitive to variation in food availability and the mean productivity of the food resources. Moreover, the magnitude of the effect of fisheries and food availability differed between cod- and herring-like life histories. These evolutionary effects are shown in Figures 3–4 as gradients in population mean *r* arising in response to changes in model conditions. Because *r* is solely genetically determined, changes in mean *r* reflect evolution. In both cases, for both cod- and herring-like fishes, resource allocation strategies evolved in response to both harvest mortality and variability in food availability described in greater detail below.

**Figure 4.**
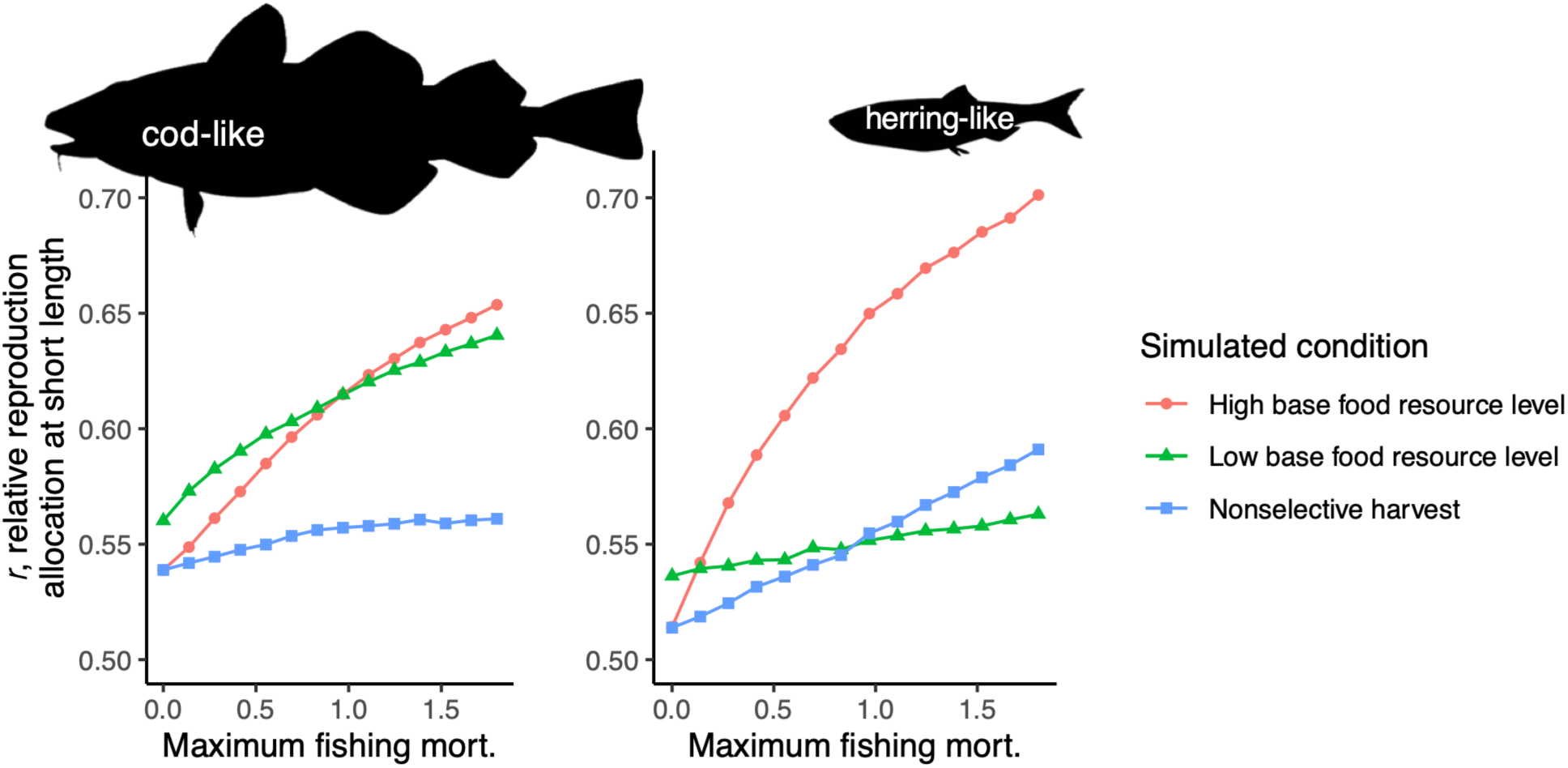
Average evolutionary change in the parameter *r* that controls how quickly fish shift from prioritizing growth to reproduction as they grow for simulated populations. Higher *r* leads to higher reproductive investment and slower growth at earlier ages/smaller sizes. Results are shown for multiple different imposed levels of harvest and two imposed levels of base food resource level (without any added random variation in food), after 40 yrs. of harvest. Results are shown for both the cod-like and herring-like models. The x axes show the value of maximum fishing mortality, F_max_. F_max_ is the maximum possible value of fishing mortality for fish that are large enough to be susceptible to fishing (Equation 5). The y axis is the average value of *r* (Equation 1a) in the population, with each point an average outcome across 500 model runs each, on a standardized scale where 1 is the maximum value possible and 0 the minimum. The value of r is determined only by genotype and by random developmental stochasticity, so change in the average value of *r* in this model implies genotypic evolution. The starting average value of *r* in the model is halfway between the possible high and low extremes (0.5 on the normalized scale for *r* in Figures 3 and 4, also see *r* in Supplemental Table 2), so values above 0.5 represent an evolutionary increase in *r*. Color (and point shape) show two different levels of initial base carrying capacity for the food resource (without any random noise added into this value). Color and point shape are also used to show a third scenario where harvest is nonselective with respect to fish length (for this scenario, base food level is at the High value of the two base food levels). As this figure is just showing different base food levels, and without random variation in *K_R_* of the food resource, there is minimal spread in the results and thus error bars would not show any spread at this scale, so they are not included.

**Figure 5.**
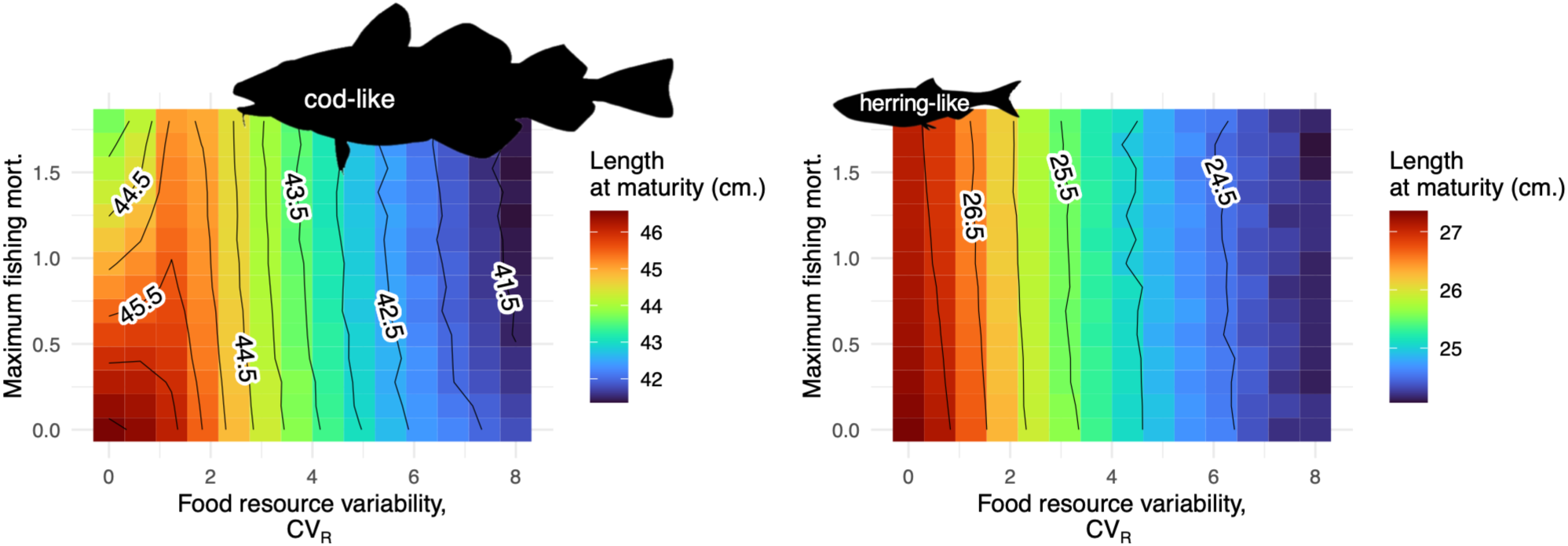
Mean length at first reproduction across different levels of harvest and food variability after 40 years of a particular harvest treatment. The x axis shows the size of the coefficient of variation in the temporally autocorrelated random values of the food resource carrying capacity, so larger x values mean more variability imposed in food resource carrying capacity. The y axis, as in Figure 3, shows the maximum fishing mortality, F_max_. F_max_ is the maximum possible value of fishing mortality for fish that are large enough to be susceptible to fishing (Equation 5). Each colored pixel (with values shown by both the colors and the contour lines) shows the mean length at first reproduction for 500 model runs after 40 years of fishing. Results are shown for both the cod-like and herring-like models.

Food variability had strong evolutionary effects on *r* in both species but with greater change in herring relative to cod (Figure 3) over the 40 years. With zero fishing mortality (when the y-axis equals zero in Figure 3), herring showed over five times as much evolutionary change as the cod due to food variability (Figure 3). As the coefficient of variation of food stochasticity (*CV_R_*) increases from 0 to 8, the standardized value of this parameter *r* for herring increases from ∼0.52 to ∼0.64, a 23% difference (Figure 3). For cod, it increases from ∼0.54 to ∼0.56, a 4% difference (Figure 3). Such evolution occurred without manipulation of the mean of the carrying capacity of the food resource, only manipulation of its stochasticity.

Unlike this food-driven evolution, harvest-induced evolution was of similar magnitude for herring- and cod-like models (Figure 3). In the case of herring, increasing the maximum length-selective fishing mortality from 0 to 1 caused an evolutionary increase in *r* by 25% when food was relatively constant (Figure 3). In the case of cod, increasing the maximum length-selective fishing mortality from 0 to 1 caused an evolutionary increase in *r* by about 22% when food was relatively constant (Figure 3).

Food variability and harvest had interactive effects on evolution in *r* in herring, but no such interactive effects were consistently observed for cod. Specifically, while increases in food variability induced genetic evolution in energy allocation (i.e. changes in parameter *r*), in herring they also suppressed evolutionary effects of fishing on energy allocation. In the herring-like model, there was an antagonistic interaction between food variability and harvest intensity where food variability reduces evolution due to harvest, for a given amount of time (40 years) in which evolution was allowed to occur. This interaction is shown in Figure 3; the change in value of *r* along the y-axis is smaller at high x-values. In Figure 3, where the x-axis is zero and the carrying capacity of food is constant, *r* for herring is about 0.52 when maximum fishing mortality is zero at the low end of the y-axis. When the food carrying capacity is constant and the maximum fishing mortality on the y-axis is 1, the value of *r* as shown by the colors and contours is about 0.65, 25% higher than at fishing mortality of zero. But when food resource variability was increased eight-fold (*CV_R_*=8), the evolutionary increase in *r* in response to fishing mortality of 1 was much smaller, less than 5%. At *CV_R_*=8, the *r* value for herring under no fishing is 0.64, and it is 0.67 under maximum fishing mortality rate of 1 unit. However, for the cod-like model, the change of *r* in Figure 3 for the cod-like panel as fishing increases does not depend on the value of *CV_R_*.

The above results assume a consistent mean carrying capacity (*K_R_*) of base food productivity across all simulations. Simulated changes in *K_R_* also caused evolutionary changes of their own. Decreasing *K_R_* by 80% induced evolution of higher *r* and thus a faster life-history strategy for herring (but not cod) and the magnitude of this effect increased with harvest (Figure 4). In the absence of harvesting (left side of panels in Figure 4), the evolutionary response to decreased base food productivity is small. Specifically, for either taxon, there is less than 5% difference between the response (*r*) under the low base food level and high base food level. For cod, *r* is ∼ 0.56 vs. ∼ 0.54 when base food levels are low vs high, respectively; for herring they are ∼ 0.53 and 0.51. However, for herring the increase in *r* diverges among base food level (*K_R_*) scenarios as harvest increases, but does not for cod (Figure 4). Decreasing base food productivity reduces the slope of the evolutionary response of *r* to fishing by more than 90% in herring with only minimal reductions for cod (Figure 4). Specifically, under low base food levels (the green line with triangles in Figure 4), increasing *F_max_* in the herring-like model from 0 to 1.8 increases *r* from around 0.54 to just 0.55 under low base food levels, but from 0.51 to 0.7 under high base food levels. In contrast, in the cod-like model, increasing fishing mortality from 0 to 1.8 leads to an increase in *r* from 0.56 to 0.64 under low food vs. 0.54 to 0.65 under high food. Maximum yield of the fishery across different levels of fishing mortality was lower for both cod and herring under consistently low food (Supplemental Figures 1-2, Supplemental Results). Total amount of simulated absolute time in years affected the amount of evolution occurring across both species, under either low or high food (Supplemental Figure 3).

The evolutionary change in *r* due to harvest mortality also depends on whether the fishing pressure is selective of large body lengths (Figure 4). When harvest is not selective in this way, the amount of evolutionary change in *r* due to fishing is much smaller, although the importance of selectivity for this evolutionary effect is smaller in the herring-like model. Under nonselective harvest with the high base food levels regime, shown on the figure by blue squares, increasing harvest caused *r* to go from 0.54 just to ∼0.55 in the cod-like model. This is ∼10% of the magnitude of harvest-induced evolution’s effect when under the selective harvest condition. In herring, which attain much shorter maximum lengths than cod, nonselective harvest at a maximum fishing mortality of 1.8 causes *r* to go from around 0.51 to 0.59.

### Plastic effects of food and harvest and their relationship to evolutionary effects

Food variability, base food productivity, and fishing pressure had plastic, non-evolutionary effects on growth and maturation in some cases. Lower base food productivity or higher food stochasticity generally reduced structural growth (i.e. smaller length-at-age) and length at maturity (Figure 5). Length at maturity declined by more than 5% in both species as the coefficient of variation for food increased from *CV_R_*=0 to *CV_R_*=8 (under no fishing). These outcomes resulted from the plastic response of undernourished fish having limited resources for continued growth in length. We know that these outcomes resulted from plasticity because they persisted when heritability was removed – i.e., evolution was “turned off” – in the *r* trait (Supplemental Figure 4). Specifically, as food variability increased from zero to the highest level, length at maturity for cod declined from 46-47 cm to below 42 cm. This effect is shown by change in length at maturity (shown by the color of the pixel, and with contour lines) along the x-axis in Figure 5 and Supplemental Figure 4.

Fishing impacts on length resulting from plasticity were largely negligible (Supplemental Figure 4). Only when base food levels were comparatively very low (Supplemental Table 3) was there a small effect. Specifically, we can observe such a plastic effect when base food level is reduced 80% from the higher base food level scenario, and when heritability in *r* is removed and thus we allow no evolution. Under the low base food level scenario, increasing fishing mortality from 0 to 1.8 leads to only slight increases (by 5mm or less, 1% or less of body length) in length-fat-age and length at maturity with greater fishing pressure in the absence of heritability, in the cod-like model (Supplemental Table 3).

Evolution in length-specific reproductive investment (i.e., in *r*) resulting from high harvest did not necessarily cause phenotypic change in maturation, even with genotypic changes. For cod, plasticity in maturation age when food was variable obscured effects of evolution in *r* on maturation length. In cod, length at maturity declines by 6% with higher fishing but only at low levels of food variability (Figure 5). This decline is illustrated in Figure 5 when food variability is low. Length at maturity declines with fishing mortality from over 46.5 cm to below 44 cm (Figure 5), a change due to evolution in *r.* But for higher levels of food variability there is much less change with harvest, less than a 1% reduction going from no to highest harvest (Figure 5) at the high end of the x-axis, where there is little or no decrease in the colors/contours with increasing y values. The fish in the cod-like model mature at average ages younger than three years when food is constant, but at average age at or above 3 years at intermediate-to-high levels of food variability (Supplemental Figure 5). This delaying maturation in response to low or variable food occurs plastically without evolution (Supplemental Table 3). An increase in length at maturity under increased food variability specifically for the most heavily fished cod is because the cod in our model at 3 years of age or older are still longer than age-2 fish, because of their older age, even with intermediate levels of food stochasticity slowing their growth. We can see this increase in Figure 5 by, for heavily harvested fish (high y-axis value), comparing length at maturity under *CV_R_* of 2-3 vs. under constant food where variability is 0. When maturing at older ages, the fishing-induced variation due to evolution in *r* does not have as much of an effect on the phenotype of length at maturity.

In herring, we observed only very slight declines in length at maturity due to harvest-induced evolution. This effect is very small, a change of 5mm. or less with an increase from 0 to 1.8 units of fishing mortality, at any level of base food productivity or food variability (Figure 5).

This small change was despite the fact that fishing caused evolution leading to higher *r* and more length-specific reproductive investment (Figure 3).

### Direct and indirect effects of temperature on fish and interactions with harvest induced evolution and plasticity

Temperature variability had no consistent effects on evolution in the fish in our model (Supplemental Figure 6). For the cod-like model, at a maximum fishing mortality in Supplemental Figure 6 of 1.0, the value of the genetically varying physiological parameter *r* stayed between 0.62-0.63 across the temperature variability values simulated. The effect of temperature variability on evolution in the herring-like model is even smaller (Supplemental Figure 6). In addition to having no effect on its own, temperature variability does not significantly change how *r* evolves in response to fishing, i.e., temperature variability does not have an interactive effect with fishing on *r*. Consequently, all of the phenotypic variation of our modeled fishes in response to temperature is due to their plasticity in response to temperature altering both their physiology and their food resource.

The direct physiological effects of temperature explain some of the variation in our modeled fishes’ growth in response to temperature variability. However, the indirect effects of temperature on fish via effects on their prey populations can quantitatively and qualitatively change these growth outcomes. In panels e and f of Figure 6, showing results of length-at-age for the cod-like model across temperature variability levels on the x-axis, we see an increase in length-at-age with higher temperature variability whether or not the fishes’ food source is affected positively or negatively by temperature. However, the magnitude of this increase in length-at-age is different. When the food source for the cod is negatively affected by higher temperatures, increasing temperature variability causes length at age 3 to increase only from about 46.6 cm. to about 47.6 cm. for fish at a fishing mortality on the y-axis of 1.0 (Figure 6f).

**Figure 6.**
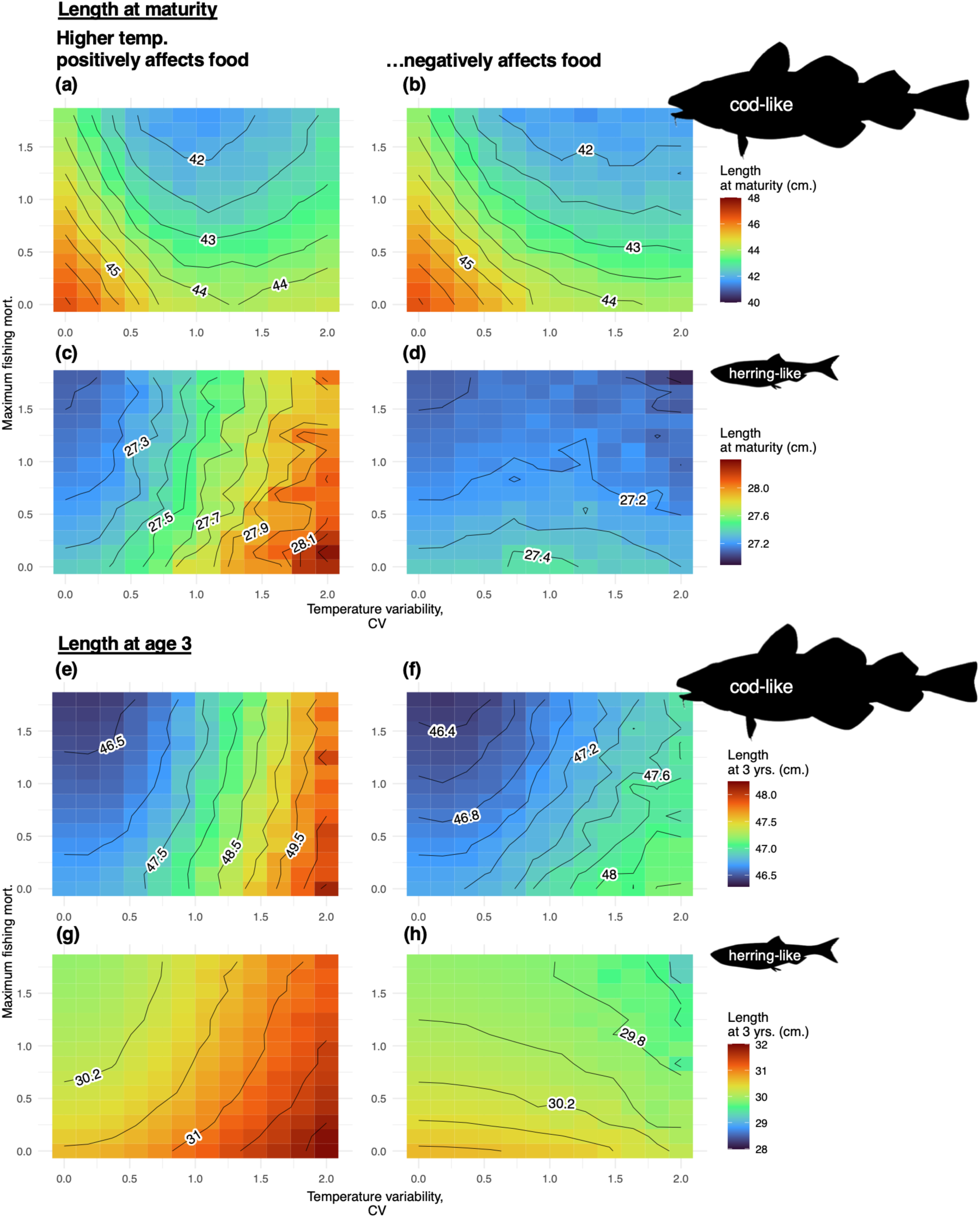
Variation in length at maturity, and length at age 3, across different levels of harvest and temperature variability. Variability in the externally imposed temperature variable affects multiple parameters of the biology of both simulated fish and their food source. (a-d) These panels show the variation in length at maturity with different levels of maximum harvesting mortality and levels of stochasticity in the temperature. The x axes show the size of the coefficient of variation in the temporally autocorrelated random values of temperature, so larger x values mean more variability imposed in temperature. The y axis, as in Figure 3, shows the maximum fishing mortality, F_max_. F_max_ is the maximum possible value of fishing mortality for fish that are large enough to be susceptible to fishing (Equation 5). Each colored pixel (with values shown by both the colors and the contour lines) shows the mean length at first reproduction for 500 model runs after 40 years of fishing. Results are shown for both the cod-like and herring-like models. (e-h) These panels show the variation in length at 3 years of age with different levels of maximum harvesting mortality and levels of stochasticity in the temperature. The x axes show the size of the coefficient of variation in the temporally autocorrelated random values of temperature, so larger x values mean more variability imposed in temperature. The y axis, as in Figure 3, shows the maximum fishing mortality. This value F_max_is the value of fishing mortality for large fish that are in the size range receiving 100% of maximum fishing mortality (Equation 5). Each colored pixel (with values shown by both the colors and the contour lines) shows the mean length at age 3 yrs. for 500 model runs after 40 years of fishing. Results are shown for both the cod-like and herring-like models. Fish silhouettes courtesy of phylopic.org (CC0 1.0 Universal Public Domain Dedication).

However, if the temperature is positively affecting their food source, the cod vary instead from about 46.6 to over 49.5 cm. over this range of variation in temperature variability values (Figure 6e). In the herring-like model, the fishes’ food source being negatively affected by higher temperatures completely erases any trend of fish length-at-age increasing with temperature variability. Figure 6h shows that herring length-at-age decreases by about 0.2cm. going across the range of temperature variability values simulated at a given fishing mortality value on the y-axis. By contrast, in Figure 6g where the temperature effects on the herrings’ food source are positive, herring length-at-age increases with food variability by over 1.2 cm.

In addition to affecting the growth in body length of cod, temperature stochasticity also affects at what lengths and ages cod have acquired enough reversible mass to be able to initiate maturation and reproduction. Supplemental Figure 7 shows that cod mature at younger ages at higher levels of temperature variability. The cod mature at as young as 2.4 years old at the highest levels of temperature variability, as compared to 3 years of age under a constant environment and no fishing. Figure 6a and b show variation in length at maturity for the cod across different fishing levels and different levels of temperature variability. These panels show changes in cod length at maturity from simultaneous effects of increased growth and accelerated maturation due to temperature variability. Under both of the scenarios in Figure 6a, where higher temperature is positively affecting the cod’s food source, and Figure 6b, where temperature is negatively affecting the cod’s food source, length at maturity initially decreases by at least 2cm with an increase from no temperature stochasticity to temperature stochasticity with a CV of 1.0 along the x-axis. Length at maturity decreases because the fish are maturing earlier, even as their rate of growth in length is increasing with increasing temperature variability. However, at even higher levels of temperature stochasticity, the effects of such temperature spikes on growth in length may dominate over this effect of a reduced age at maturity in determining what the length at maturity is. We see this positive effect on growth in length come through in Figure 6a, where at the highest levels of temperature stochasticity, temperature stochasticity increases length at maturity. In Figure 6b, the food source of the cod is negatively affected by high temperature, and consequently the increases in a cod’s growth in length with increasing temperature stochasticity are smaller than they would be if their food source were also positively affected by temperature (Figure 6e-f). Consequently, for Figure 6b, length at maturity never increases along the x-axis with increasing temperature stochasticity, it only decreases.

While variability in the food source for the cod-like model caused plastic effects that obscured fisheries-induced evolution’s effects on the phenotype of length at maturity (Figure 5), variability in temperature does not necessarily have the same effects that variability in food does, of preventing length at maturity’s change with fishing. In Figure 6b, for instance, the cod length at maturity decreases by at least 2.5cm., going from no fishing to maximum fishing mortality of 1.8, across all different levels of temperature variability (all x-axis values). In Figure 6a, where the higher temperatures positively affect the cod’s food source, the decrease along the y-axis in the length at maturity of cod is about 3cm. but goes to less than 2cm. at our highest simulated level of temperature variability–at the high end of the y-axis, the change in the outcome along the x-axis is slightly smaller. The strong effects of temperature stochasticity on growth are contributing to a plastic effect of diminishing how fisheries-induced evolution affects length at maturity.

In the herring-like model, the effects of temperature variability on length at maturity are even more dependent than they are in the cod-like model on whether the fishes’ food source is being positively or negative affected by higher temperature. Length at maturity in the herring-like model is essentially determined by body length of the fish at a given age, since the age at maturity does not vary in the herring-like model – our simulated fish in the herring-like model always mature at 1 yrs. old. Given this inherently maximally accelerated maturation from the model parameterization, the patterns of length at maturity in herring across different temperature variability scenarios (Figure 6c-d) mostly follow the pattern of herring length-at-age across different temperature variability scenarios (Figure 6g-h, discussed above). For a given fishing mortality value on the y-axis, length at maturity for herring where their food source is negatively affected by high temperature fluctuates within about 0.1cm. across different temperature variability values but shows no consistent trend (Figure 6d). In Figure 6c, we see that length at maturity increases by as much as a centimeter or more across the gradient of temperature variability for herring with a food source experiencing positive effects of temperature variability.

## Discussion

How top-down and bottom-up forces interact to shape evolution and plasticity in life history remains a central question in population and evolutionary ecology. Top-down effects like harvest or other predation can select for slower growth rates and both greater and earlier investment in reproduction (Heino et al. 2015). In many systems these effects depend on harvest selecting against older adults and/or larger sizes (Grainger and Levine 2022). Even as such adaptive evolutionary changes generally boost population growth and numerical resilience to harvest, they may leave populations maladapted to certain environmental factors or changes, making such patterns important for management consideration (Heino et al. 2015; Edeline and Loeuille 2021). These patterns emerge in our study for two very different taxa, yet such effects also depend on the productivity and stochasticity of food availability. An interaction between evolution due to harvest and due to food may occur when these distinct factors cause selection on the same genes or gene networks. Additionally, they may interact because harvest and food availability can be causally linked. For example, harvest may indirectly affect per capita food availability in a food-limited environment by decreasing both contest and scramble competition (Edeline and Loeuille 2021). Similarly, changes in productivity can decrease negative impacts of high harvest mortality. As a result, whether fished populations are food-limited, limit their food source, and/or are limited top-down by predation when they themselves are food, can all inform optimal management (Turcotte et al. 2021; Czorlich et al. 2022).

Consumer-resource dynamics also include those between human fishers and the fished stock. In our model runs the maximum fishing mortality and the selectivity were fixed for each run, but interactions between environmental variation such as climate variation and fishing fleets’ dynamic responses to a stock can determine whether a stock persists (Lima et al. 2020). The role of fish life history traits in such interactive effects is an avenue for future research. We also recognize that stock assessment researchers are often very limited in data they can feasibly acquire, making prioritization essential (Cope et al. 2023). Debate exists as to the importance of external environmental factors like climate and prey, and their feedbacks and interactions with a focal stock, in practical stock management in fisheries (Edgar et al. 2019). Ultimately, in many examples, variation in stock dynamics can be explained by variation in partly external environmental variation ranging from a stock’s food source, climate conditions, to environment-driven changes in life history traits (Young et al. 2006; Okamoto et al. 2012; Okamoto et al. 2016; Hay et al. 2019; Sguotti et al. 2019). Our modeling work here, while simplified and theoretical, adds to this strong basis for considering how food variation and harvest individually and interactively shape trait distributions in populations, which may in turn affect some populations’ sustainability. Because both environmental and genetic changes may affect shifts in key traits, it may be critical to not only measure phenotypic changes over time, but also genomic shifts and the physiological processes that regulate such emergent properties. Moreover, managers and practitioners may be aware of potential selective pressures of harvest and factor such uncertainties into fisheries risk assessment when considering evaluation of management strategies.

Our work suggests multiple mechanisms by which variation in food interacts with fishing to shape adaptive landscapes. Gobin et al. (2018) previously identified one such mechanism in a modeling study. They suggested that low food may reduce fisheries-induced evolution because slower growth yields fewer fish and older fish entering the harvestable body size range (for a fishery with a rigid lower limit to harvestable body size). The same dynamics are likely present in our results (Supplemental Figure 2). It is difficult to disentangle the effects of different food conditions on fish generation time (which influences rate of possible adaptive evolution) from effects of different food conditions on amount of adaptive evolution. However, across the two different food conditions shown for the herring-like fish in Figure 4, herring age-at-maturity and generation time did not change. By contrast, the amount of herring being removed by the fishery was reduced dramatically under low food or variable food (Supplemental Figure 2). Therefore, this reduction of the strongly size-selective effects of selection from harvesting when food, and growth, are reduced, likely played a role for our herring-like model in low food reducing the magnitude of fishing-induced evolution in *r*.

We extended Gobin et al.’s approach by allowing our fish population to evolve in response to food variation itself, and by allowing our fish population’s food resource to dynamically vary in response to fish consumption. Gobin et al.’s model included negative effects of low food on growth; in our model, we also allow food deprivation to have direct mechanistic effects on mortality and reproduction. We show that low or variable food on its own can cause evolution of faster life histories and smaller maturation when fishing is absent. However, variable food can also reduce the amount by which reproductive investment evolves in response to a given amount of fishing pressure. These results indicating that variability in food alone is sufficient to drive evolution of life history is consistent with theory that variable adult survival or risk of not enough resources to reproduce should favor evolution of more reproduction (Darwin 1859; Schaffer 1974; Stearns 1983).

In addition to food availability’s direct evolutionary effects (Diopere et al. 2018), food availability may interact with harvest’s effects on traits because of how harvest indirectly may shape fishes’ food acquisition (Taylor and McIlwain 2010). When prey productivity is low and the fish limit their prey resource, fishing in our cod-like model can slightly increase food intake of individual fish, increasing growth and survival. This plastic density-dependent effect has been proposed in the past as opposing and hiding evolutionary reduction in length-at-age phenotypes (Conover and Munch 2007). However, a plastic effect of relaxation of competition represents an alteration of selective pressures that by extension can affect evolution. Population density in nature often has evolutionary effects, independent of predation, with reduced density decreasing reproductive investment in some cases (Schrader and Travis 2012). However, in some cases reduced density can also select for less somatic growth in body length (Bouffet-Halle et al. 2021), when growing longer to compete more effectively is both costly and less advantageous. Additionally, in our model results, it is likely that differences in generation time can explain the food-scarcity-driven reduction in fishing-induced evolution in Figure 4. Under either highly variable or constantly low food conditions, age at maturity of the cod-like fish in our model was increased (Supplemental Figure 5). Longer generation time and fewer generations per unit time is expected to reduce the amount by which a population can, in a given amount of time, undergo a given amount of evolutionary response to selection (Orive et al. 2017). Rates of evolution in life history traits in harvested or predated populations ultimately depend on the values of those life history traits themselves, because life history traits influence generation time and therefore rate of evolution (Devine et al. 2012; Osmond et al. 2017).

Plastic effects of the environment on phenotypes can obscure the emergent phenotypic effects of a given genotypic change on some phenotype (Sultan 1987). For our cod-like model, phenotypic effects of genetic responses to fishing can be suppressed by high food variability plastically delaying maturation. For herring, genetic evolution in length-dependent resource allocation has minimal effects on length at maturation and none on age at maturation at almost *any* food level. Aspects of how the herring model is parameterized force rapid maturation in the first year of life of this rapid-life-history taxon, even when evolution is changing its reproductive investment. For simplicity and generality, we do not attempt to capture the full suite of environmental and physiological interactions underlying maturation (Pauly 2021a). How best to represent the link between fish maturity and underlying physiology in models is still a subject warranting further empirical research. However, the stasis we see in maturation despite substantial physiological evolution also highlights the important fact that the absence of observed phenotypic change in a population does not imply absence of evolution. Change may occur in physiological phenotypes which are more difficult to measure in the field than age and length (Hollins et al. 2018), or plasticity may counter evolution, keeping phenotype constant (Potter et al. 2021). Harvested fish are very biodiverse, and variation relevant to eco-evolutionary life history shifts in harvested fish may go beyond the traits we simulate in this model. For some taxa, traits mediating harvest-induced eco-evolutionary dynamics may include more unique traits like protogyny in many reef fish fishes (Sattar et al. 2008), social learning (MacCall et al. 2019), ontogenetic niche shifts (Huss et al. 2012), or extreme metabolic plasticity (Spindel et al. 2020).

Our work supports the possibility that some trait distribution changes appearing to be evolution in fished stocks may be plastic and/or a response to non-harvest factors. Simultaneous reduction in both length and age at maturity has been argued to be evidence of evolution in fish. However, empirical evidence suggests simultaneous reduction in both age and length at maturity can occur plastically (Uusi-Heikkilä et al. 2011), and our model likewise supports an expansive view of what combinations of change in age and length in maturity can be plastic. According to our cod model (in Figure 5 and Supplemental Figure 5, for cod maturing in 3 yrs. vs. those maturing younger), reducing food stochasticity can sometimes plastically reduce maturation length and age. Going from some to no food stochasticity, at high fishing levels, we see simultaneous decreases in age and length at maturity without seeing corresponding evolutionary change. Reznick (1993) described that seasonally breeding fish that can only breed at one time of year (as in our model) may show a positive correlation between likely ages and lengths at maturation, because they may delay maturation for a long time if they skip first spawning.

However, some life history theories have suggested that adaptive plasticity in life history should not allow plastic decreases in both length and age at maturity (Day and Rowe 2002), which may have led to incorrect inferences of harvest-induced evolution (Hutchings and Kuparinen 2020).

Both our work and prior theoretical research suggests evolution in traits like length at maturity in response to selective harvest may be relatively small. This expectation may aid in inferring which life history trait changes in harvested populations may result from harvest-induced evolution, plastic, and/or a response to non-harvest factors. Andersen and Brander (2009) used a model like ours which, like ours, accounts for some energetic and fecundity tradeoffs of growing less in length. They showed evolutionary decline in maturation length of ∼0.1% per year due to fishing. This change is consistent with our evolutionary decline in cod length at maturity by ∼6% over 40 years under constant food (left of cod panel in Figure 5). This rate is similar to that identified in an evolutionary modeling study by Forestier et al. (2020).

Where larger changes in life history are observed in fish, these may be in part due to plastic responses to fishing, or factors other than fishing. Plasticity generally does not accelerate life history due to harvest in our model, but some fish may experience plastic responses to harvest unaccounted for in our model, such as response to release from intense size-based contest competition, or other forms of anticipatory plasticity (Bouffet-Halle et al. 2021; Edeline and Loeuille 2021; Harmon et al. 2023). Predicting potential fisheries-induced evolution requires more empirical work on what patterns of trait variation in fish are heritable, amounts of heritable and non-heritable evolution in traits, and traits’ genomic basis and genetic correlations (Irwin and Carter 2013; Enberg and Jørgensen 2017; Therkildsen et al. 2019).

Of environmental factors beyond harvest, anthropogenic climate warming is one of the most significant affecting harvested fishes (Cheung et al. 2013). Our work shows that changing temperature can alter how fish population phenotype distributions respond to harvest. Further, we show this interaction may depend on how a focal population’s food source is affected by temperature. Jessen et al. (2025) used an individual-based model to study effects of increasing temperature on fisheries-induced evolution in cod. Their model differed from ours in some bioenergetic details; in which traits were made genetically variable and evolvable; and in that they modeled temperature effects solely on focal organisms, not indirect effects through food sources. Their paper and our work agree that, in some cases, increases in temperature may cause increased growth rates in fish, and suppression of change in phenotypic distributions due to harvest. Both our paper and theirs leave open the question of how these patterns may further change in populations where higher temperatures lead to decreases, not increases, in aerobic scope and/or total surplus energy availability. The importance and even direction of temperature effects on oxygen and energy availability vary between and within fish taxa (Holt and Jørgensen 2015; Audzijonyte et al. 2019; Wootton et al. 2022). And more work is needed to understand the complex outcomes for species interactions when interacting species simultaneously experience climate effects (Dee et al. 2020), and how to link individual-level temperature dependency and population-level outcomes (Rezende and Bozinovic 2019; Munch et al. 2023).

Human harvest has shaped evolutionary pressures in harvested taxa for millennia (O’Dea et al. 2014). The intensity of anthropogenic environmental impacts makes rapid evolution an important force to consider in ecology and conservation even as much rapid trait change may not be evolutionary (Sanderson et al. 2022). Here, we demonstrate that the evolutionary effects of harvest on life history traits may be prevented, masked, or enhanced as food resources change in productivity or variability. Recent calls have been made to incorporate more trait distribution data and investigation of evolution into management of harvested populations. Yet the practical importance for, and capacity of, management to collect and use trait data likely varies dramatically across fisheries (Barnett et al. 2019; Hutchings and Kuparinen 2021). Our research suggests that even where harvest has plastic or evolutionary effects on traits, these are heavily modulated by food variation. Further work is needed to determine how these processes play out in experimental and wild populations. Moreover, climate change continues to impose abiotic changes such as warmer oceans and lower oxygen concentrations that directly reduce energy availability for fish, and these may determine surplus energy in some fish as much or more than prey abundance (Audzijonyte et al. 2020; Pauly 2021b). Regardless of the challenges and uncertainties, consideration of consumer-resource dynamics is likely to be crucial in any effective implementation of trait-based approaches to manage and understand wild populations.

## Supporting information

Supplemental Word doc, with supplemental figures, tables, and info

Code and simulated data files

## Notes

### Competing Interest Statement

The authors have declared no competing interest.

