## Supplemental Word doc, with supplemental figures, tables, and info for "Environmental variability alters and obscures fisheries-induced evolution"

**Supplemental content for “Environmental variability alters and obscures fisheries-induced evolution”**

Supplemental Figures 1-7 (included below)

Supplemental Methods and Results

Supplemental Table 1 (included below)

Supplemental Table 2 (included below)

References for Supplemental Tables 1-2

Supplemental Table 3 (in separate Excel file)

Technical Supplement I (hosted here): https://ecoevoenergymodels.shinyapps.io/Supplement/

Link to simulated data and code at an anonymized GitHub repository (also linked to in the above shinyapps site): https://anonymous.4open.science/r/ecoevoharvestmodel-420C. The large data files are also included in a zipped folder supplementally as part of the submission.

A link to the GitHub repository with the simulated data and code in a public repository (Zenodo) will be provided if the paper proceeds in the review process.

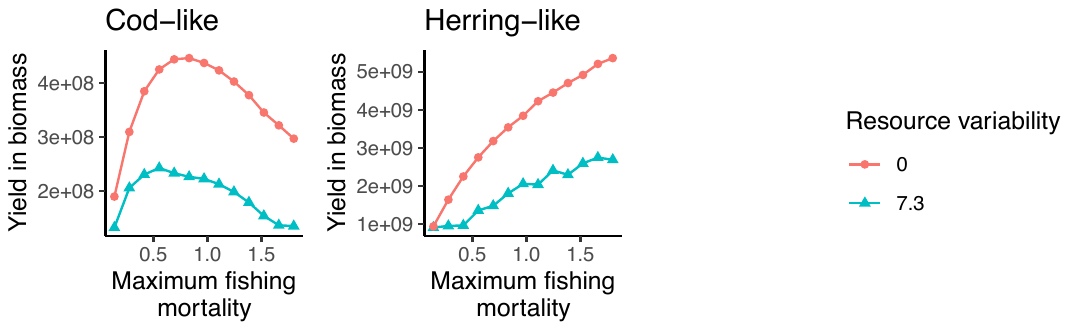

Supplemental Figure 1. Mean per-day yield from harvest in units of total biomass, at different levels of food variability and maximum fishing mortality. The explanatory variables shown here by the x axis and by the point colors/shapes are analogous to the maximum fishing mortality and resource carrying capacity random variation variables from the main text Figures 3 and 5.

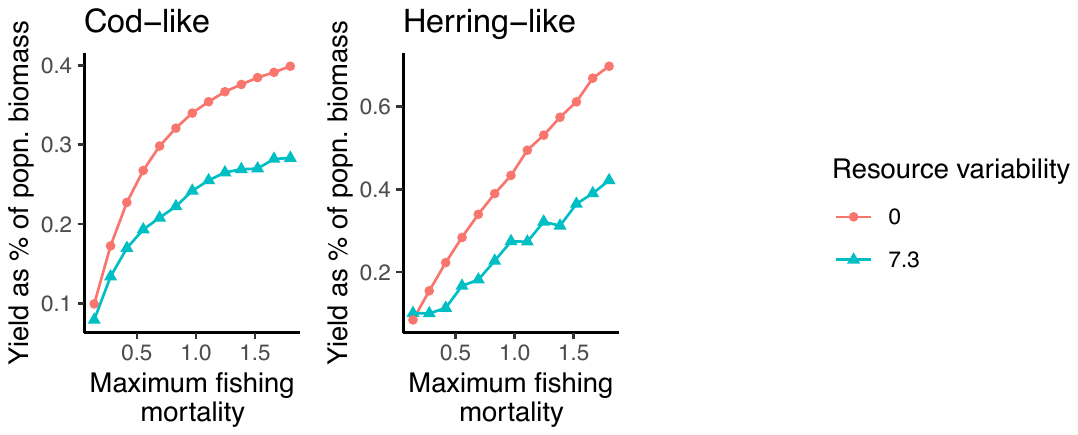

Supplemental Figure 2. Mean per-day yield from harvest, with the yield given as the percentage of the total population biomass, at different levels of food variability and maximum fishing mortality. The explanatory variables shown here by the x axis and by the point colors/shapes are analogous to the maximum fishing mortality and resource carrying capacity random variation variables from the main text Figures 3 and 5.

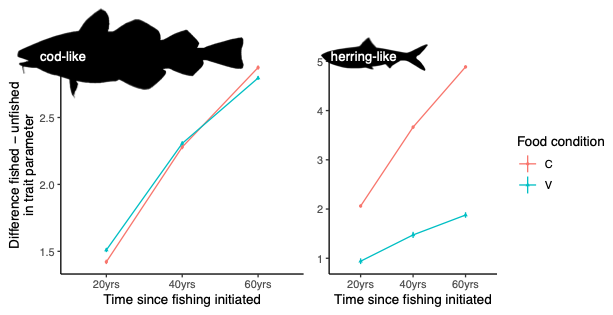

Supplemental Figure 3. The difference in the value of the genetically-controlled heritable physiological parameter *r*, between treatments with no fishing and treatments with maximal (maximum instantaneous fishing mortality of 1.8 units) fishing. This difference is shown across different amounts of time since introducing fishing. The y-axis represents the comparison in the mean of *r* for 500 model runs with 1.8 units’ maximum fishing mortality, and no fishing mortality, for each of the three simulation lengths (20, 40, or 60 yrs.). The time on the x-axis does not include the amount of time the models were run for equilibration of population dynamics before fishing either was or was not introduced (80 years). The comparison here between fished and unfished populations was repeated, both with no stochastic variability in food carrying capacity, and high stochasticity in the food carrying capacity (coefficient of variation of 2). These two food conditions are shown in the two different line colors and point shapes. Standard deviations of sample mean differences between 500 fished and 500 unfished model runs are shown for each combination of time horizon and food condition in vertical error bars, but, on this scale, these are only large enough to be visible for the herring in the highly variable food condition.

Supplemental Figure 4. Cod population mean length at first reproduction, after 40 years of a particular harvest treatment, across different combinations of harvest and food variability. The difference between this figure and the main text’s Figure 5 for cod is that this figure does not allow evolutionary processes to occur. These are results from the cod-like model but without any genetic diversity in the fish populations (consequently, no evolution is possible). Each square pixel is a mean length at first reproduction for 500 model runs. The x axis shows the size of the coefficient of variation in the temporally autocorrelated random values of the food resource carrying capacity. The y axis shows the maximum fishing mortality. This maximum fishing mortality is the value of fishing mortality for large fish that are in the size range receiving 100% of maximum fishing mortality (Equation 5).

Supplemental Figure 5. Cod population mean age at first reproduction, after 40 years of a particular harvest treatment, across different levels of harvest and food variability. Each square pixel is a mean age at first reproduction for 500 model runs. These are results from the cod-like model, *with* evolution occurring (distinct from Supplemental Figure 4). This figure shows the mean ages for the model runs for which the mean lengths are shown in the cod panel of Figure 5 – but here just showing age, rather than length. The x axis shows the size of the coefficient of variation in the temporally autocorrelated random values of the food resource carrying capacity. The y axis shows the maximum fishing mortality. This maximum fishing mortality is the value of fishing mortality for large fish that are in the size range receiving 100% of maximum fishing mortality (Equation 5).

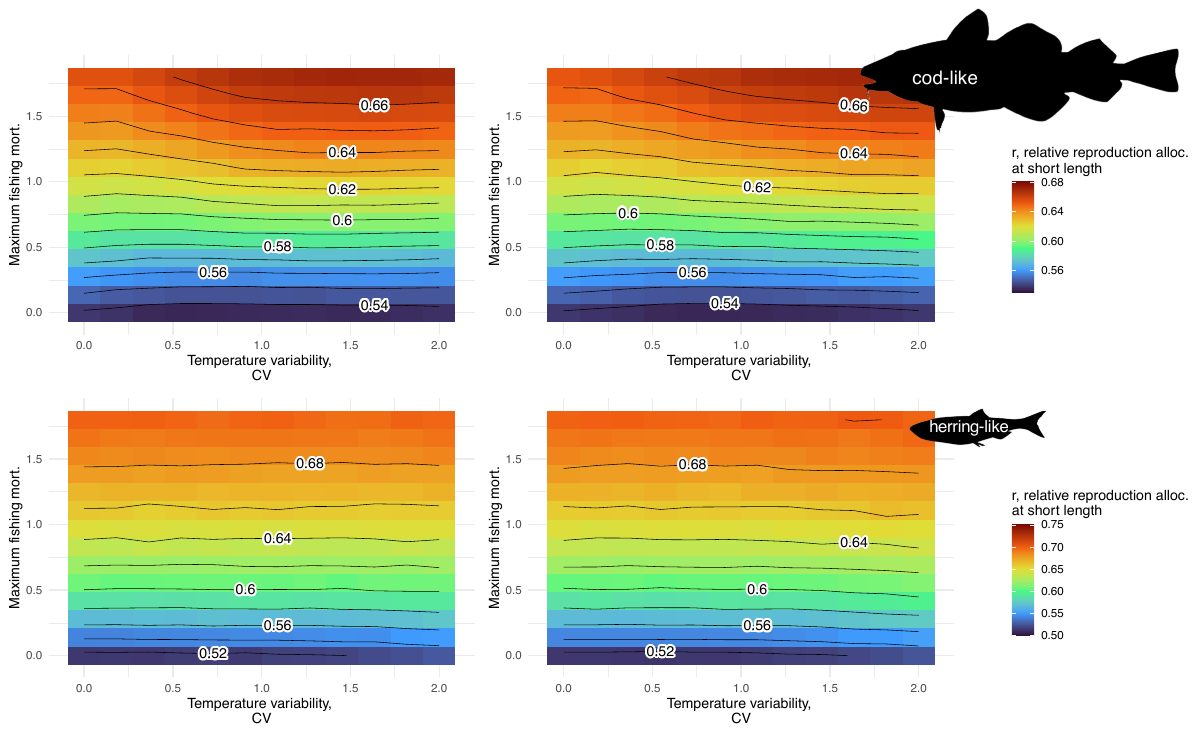

Supplemental Figure 6. Average evolutionary change (driven only by evolution in populations’ breeding values) of simulated populations under different levels of harvest and temperature variability, after 40 yrs. of harvest. The outcome variable shown by the contours and the colors of the pixels is the average value of *r* (Equation 3c) in the population, averaged across 500 model runs each for each x-y combination, on a standardized scale where 1 is the maximum value possible and 0 the minimum. The x axis shows the size of the coefficient of variation in the temporally autocorrelated random values of the temperature, so larger x values mean more variability imposed in temperature. The y axis shows the value of maximum fishing mortality. This maximum fishing mortality $F_{\max}$is the maximum possible value of fishing mortality for fish that are large enough to be susceptible to fishing (Equation 5). The value of *r* is determined only by genotype and by random developmental stochasticity, so change in the average value of *r* in this model implies genotypic evolution. The starting average value of *r* in the model is halfway between the possible high and low extremes (0.5 on the normalized scale for *r* in Figures 3 and 4, also see *r* in Table 2), so values above 0.5 represent an evolutionary increase in *r*. Results are shown for both the cod-like and herring-like models.

Supplemental Figure 7. The cod-like population mean age at first reproduction, after 40 years of a particular harvest treatment, across different levels of harvest and temperature variability. Panel (a) shows the results for model runs where temperature positively affects the population growth of the fish’s food resource, and panel (b) shows results from scenarios where temperature negatively impacts the fish’s food. Each square pixel is a mean age at first reproduction for 500 model runs. This figure shows the mean ages for the model runs for which the mean lengths at maturity are shown in the cod panels of the main text’s Figure 6 – but here just showing age, rather than length at maturity. The x axis shows the size of the coefficient of variation in the temporally autocorrelated random values of the food resource carrying capacity. The y axis shows the maximum fishing mortality. This maximum fishing mortality $F_{\max}$is the maximum possible value of fishing mortality for fish that are large enough to be susceptible to fishing (Equation 5). Results for the herring-like model are not shown here because age at maturity of herring did not vary across model runs.

**Supplemental Methods and Results**

*Timescales and numbers of model runs*

We chose a 40-year horizon for model runs, following 60-year equilibration periods prior to introduction of fishing mortality levels. Any longer timescales are likely to experience long-period trends, other shocks, and ecological interactions not included in our model. Because evolution had not always reached an equilibrium by the end of this time, we also evaluated some cases for longer or shorter periods to assess any dependence on time horizon of our main results. When modeling temperature effects, we simulated at least 700 model runs for each combination of temperature stochasticity and maximum fishing mortality due to increased outcome variation. This was in contrast to 500 models runs for each combination of parameters when we were directly varying the fishes’ food source, as described in the main text.

*Notes on population dynamics of simulated fish populations*

The shifts in growth and maturation resulting from variability in food availability also affected productivity of the fish stock. Specifically, higher variability in food decreases total harvest from the population (Supplemental Figure 1). Moreover, stochastic food variability also reduces yield relative to the population biomass (Supplemental Figure 2). Yield per unit population biomass decreases by more than 25% for both taxa when resource variability is at its highest level (Supplemental Figure 2). This decrease in yield per unit population biomass results because there are fewer fish in the harvestable size range. Specifically, higher food variability leads to plastic declines in growth leading to a slower rate of recruitment into the fishery that is rigid in both its selectivity ogive and minimum size.

*Sensitivity of model outcomes to other parameters*

In the main text, we show that the response of life history traits to fishing exhibits strong sensitivity to base life history (i.e., “herring” vs. “cod”), magnitude of base food productivity, food variability, and intensity and selectivity of harvest. These dependencies include not only the magnitude and direction but whether they result from evolutionary and/or plastic responses. We also assessed sensitivity of our model results to variation in other model parameters, and these had little qualitative effect on model outcomes. We tested the sensitivity of model results to variation in heritability by reducing the narrow-sense heritability which is the proportion of total trait variance that results from additive genetic variance. We achieved this aim by increasing developmental stochasticity ($\sigma_{devo}$ in Table 1) which is the random intrinsic variation in a trait for a given genotype, where in our case the trait of focus is *r* which was the genetically varying resource allocation parameter in our models. Such intrinsic variation does not include variation caused by the external environment. Increasing developmental stochasticity increased variation in non-heritable trait values and thereby decreased heritability of *r*. However, such a reduction in heritability had essentially no effect on evolution occurring in this trait or in emergent traits after periods of 40 years of simulated evolution (Supplemental Table 3). Our base models also assume the only heritable trait is the physiological energy allocation parameter *r*. Thus, we also assessed sensitivity of results to allowing genetic variation in other traits, namely maintenance costs and length-specific food intake. These scenarios still resulted in evolution, at higher fishing, of trait values that would give smaller length at age and length at maturity. Both cod and herring-like models showed evolution of at least slightly higher maintenance costs and slightly lower food intake when selective fishing was high, resulting in smaller length-at-age (Supplemental Table 3). Reducing mortality rates from non-harvest predation and from low condition by an order of magnitude had little effect, and no qualitative effect, on evolutionary and phenotypic outcomes (Supplemental Table 3). The main effect of lowering condition-dependent mortality was to slightly accelerate evolutionary change in *r* due to harvest.

The evolutionary response of life history traits to fishing varied substantially depending on the time horizon used in our simulations. The amount of evolutionary change in response to a given amount of fishing mortality, compared to a scenario with no fishing, continued to increase if the simulations were run for 60 instead of the usual 40 years for our simulations (Supplemental Figure 3). However, the dependency of this effect on food resources, seen in Figure 3, for the herring-like model, was very robust to the choice of time horizon (Supplemental Figure 3).

|  | Will this affect **length-specific rules** setting the possible use and intake of energy? | | | | | | Will this affect **age and size at maturity**? | | | | | |
| --- | --- | --- | --- | --- | --- | --- | --- | --- | --- | --- | --- | --- |
|  | Hypothesized ***plastic*** effects? | Biological rationale | Supporting evidence | Hypothesized ***evolutionary*** effects? | Biological rationale | Supporting evidence | Hypothesized ***plastic*** effects? | Biological rationale | Supporting evidence | Hypothesized ***evolutionary*** effects? | Biological rationale | Supporting evidence |
| High food variability | **None.** Length-specific patterns of use and allocation of energy are set by parameter values of bioenergetic functions, which take body size and characteristic as inputs and give energy use as outputs. These parameters will not change plastically in this model. | Bioenergetic functions determine how fish in our model can plastically respond to the environment, but do not themselves plastically change. All phenotypic plasticity has some physiological limit set by some genetically fixed physiological factors (1). We treat all specified physiological rates as fixed at early age by just genetics and development-tal noise, though plasticity in such rates can occur (6). | 1 and 2 for review and theory of reaction norms (environ-mental variation within genetically set boundar-ies), 3 and 4 for models built this way, 5 for empirical example of strong genetic basis of physiologi-cal rates, 6 for an empirical example of energetic rates which *do* change plastically | Will select for energy expenditure for more eggs at smaller body lengths, will select for shorter lengths for a given age | Lower adult survival and greater likelihood of missed reproduction selecting for fast reproduction | 3, 7, 8 for theory, 9 for wild example | Will cause delayed maturation | Relatively high chance of insufficient resources to reproduce at a given length and age (16) | Supported by fish empirical work (16), also thresholds concept of maturation (17) | Evolutionary effects on age and size at maturity will only occur via evolutionary changes in bioenergetic functions which have bottom-up effects on maturation | In our model, maturation phenotypes don’t evolve, but underlying bioenerget-ic traits which control them may evolve. See discussion in left-most columns of rationale for physiologic-al traits as foundation for plasticity in emergent life history phenotype | 21 and 22 present evidence that maturation is linked to body length and condition more than chronological age; 17 reviews underlying physiological thresholds of maturation; 10 describes that genes affect emergent growth patterns via underlying physiological mechanisms; 23 shows how emergent function-valued traits like size at age are genetically correlated and a product of shared, underlying mechanisms rather than being independent traits themselves |
| Harvest |  |  |  |  | Lower adult survival and reproduction opportunities *in addition to* high mortality for large fish selecting for small bodies, maturers | 10 for experimen-tal and 11 for wild example (but see 12) | May speed growth and maturation | Fewer fish competing for the same resources, more food resources per capita | More mass and eggs are empirical results of harvest (18, 19) |  |  |  |
| Food variability’s interaction with harvest effects |  |  |  | Food variability will reduce harvest’s evolutionary effects | Fish are already smaller because of periodic low food (so less likely to experience selection from large-capture fishery), and are already adapted to higher adult mortality; also, benefit of competitive release due to fishing may be higher if food is constrained | 13 for modeling example (but see 14, an experiment suggesting that comp. release selects for small fish) | High harvest may increase growth when resources are limited | When food is scarce, harvest may have large effects of competitive release and increased per-capita food | The likely mechanisms for the growth-releasing effects of harvest described in 18 and 19 are food resource limitations |  |  |  |
| Fast- vs. slow-life history taxon |  |  |  | Different taxa will differ in evolutionary response | Longer-lived, larger taxa may experience less selective pressure from food variability, but more from harvest | 15 for theory of how life histories respond differently, with fast life histories requiring evolutionary change to alter fitness under resource stress | Fast-growing taxa may undergo less plastic change in life history in response to the environment | Genetically different sets of bioenergetic parameters that lead to earlier reproduction and less iteroparity can also be associated with less flexibility to food resource patterns (15, 20) | 15 is a theoretical model, 20 a review that discusses how life histories respond to environmental fluctuation |  |  |  |

[1] Sultan and Stearns 2005, [2] Stearns and Koella 1986, [3] Audzijonyte and Richards 2018, [4] Huss et al. 2012, [5] Auer et al. 2018, [6] Spindel et al. 2020, [7] Caswell 1982, [8] Adams 1980, [9] Czorlich et al. 2022, [10] Therkildsen et al. 2019, [11] Therkildsen et al. 2013, [12] Pinsky et al. 2021, [13] Gobin et al. 2018, [14] Bouffet-Halle et al. 2021, [15] Schmid et al. 2022, [16] Reznick 1993, [17] Pauly 2021a, [18] DeMartini et al. 1993, [19] Koslow et al. 1995, [20] Planque et al. 2010, [21] Policansky 1982, [22] Uusi-Heikkilä et al. 2011, [23] Irwin and Carter 2013

**Supplemental Table 1.** Hypotheses for evolutionary and plastic effects of harvest and food, and the interactions of these effects. White columns indicate hypotheses about the model’s emergent behavior; gray columns indicate dynamics which are predetermined due to the fundamental design and simplifications of the model.

| **Parameter** | **Parameter description** | **Value(s)** | **References* and notes** |
| --- | --- | --- | --- |
| **Growth and resource flux** | | | |
| *a_max_* | Maximum longevity | 25 years (cod), 15 years (herring) |  |
| *r* | Controls steepness of increase in $\lambda$ with increasing length; see $\lambda$ and Equation 3c | 0-12 (depends on parents’ assigned genotypes and developmental noise) | The starting mean value for *r*, before evolution is allowed to occur and then observed, is 6 (0.5 on the normalized scale used in Figures 3 and 4). |
| $\bar{l}$ | Length at which $\lambda$ reaches an inflection point; see $\lambda$ and Equation 3c | 0.2 meters | Appropriate value from time course of condition with length shown in results of Audzijonyte and Richards’ (2018) model; this function was originally age-based in their paper |
| *c_1_* | Scaling parameter for obtaining structural mass (g.) from standard length (m.); see equation 3d | 5735 (herring-like), 5787 (cod-like) |  |
| *c_2_* | Exponent parameter for obtaining structural mass from standard length | 3 (cod-like), 3.125 (herring-like) |  |
| $\rho_{0}$ | Multiplier for obtaining overall daily energy intake as a function of structural mass (g./day) | 0.1 (cod-like), 0.3 (herring-like) | For herring, using data on adult daily food intake per gram body mass by Rudstam (1988) and assuming 2/3 for $\rho_{1}$ |
| $\rho_{1}$ | Exponent for obtaining overall daily energy intake as a function of structural mass | 2/3 |  |
| $\rho$ | Energy acquired before maintenance costs are subtracted and before *E:S* ratio is optimized | See Equation 1 |  |
| $\rho_{net}$ | Energy acquired after maintenance costs are subtracted and before *E:S* ratio is optimized | See Equation 3b |  |
| $\lambda_{opt}$ | The length-dependent optimal value of *E:S* (ratio of reversible to structural mass) | See Eqn. 3c |  |
| $\lambda_{max}$ | Maximum value of *E:S* (ratio of reversible to structural mass) fish will attain | 1.3 (cod-like), 1.13 (herring-like) |  |
| *c_S_* | Multiplier for obtaining cost of maintenance of structural mass (grams/day, per gram) | 0.003 (cod-like), 0.03 (herring-like) |  |
| *c_E_* | Multiplier for obtaining cost of maintenance of reversible mass (grams/day, per gram) | 0.0003 (cod-like), 0.03 (herring-like) |  |
| *k* | The sum of *c_S_S* and *c_E_E*, the instantaneous total body maintenance cost |  |  |
| *e_E_* | Efficiency of conversion of food or stored energy to reversible mass (grams/day, per gram) | 0.9 |  |
| *e_S_* | Efficiency of conversion of food or stored energy to structural mass | 1/3 |  |
| *o_0_* | Scale parameter for non-gametic cost of reproduction in grams reversible mass = function of structural mass (grams) | 6 (cod-like), 1.25 (herring-like) | For herring: assuming 15-25% mass loss at reproduction (Megrey et al. 2007), that minimum *E*/*S* at first reproduction = 0.7 (from Huss et al), and assuming (from Huss et al) that half of reproductive costs go to gametes |
| *o_1_* | Exponent parameter for cost of reproduction = function of structural mass | 0.6 |  |
| *w* | Fraction of reversible mass fish will spend on reproduction, if reproducing | 0.5 (herring-like), 0.6 (cod-like) | For herring, this value will (given the previously established cost parameter for reproduction *o_0_*) yield reproduction for a non-starved fish at 30cm. (a typical length by which clupeids have already matured, Milton et al. 1993) |
| **Mortality (see Equations 4-5)** | | | |
| *m* | Base mortality rate | 0.2 yearly instantaneous mortality |  |
| *m_Pmax_* | Maximum mortality rate including predation and base mortality, but before condition-dependent mortality | 0.5 |  |
| *m_Cmax_* | Maximum amount added to the base mortality rate due to low (essentially zero) condition | 18 |  |
| *z_c_* | Parameter setting the decline in condition-dependent mortality as condition improves | 7 (cod-like), 35 (herring-like) | For herring, a 5x faster decline in this form of mortality as condition improves (i.e., strong starvation mortality only operates on herring at *very* poor condition, compared to cod) results in negligible starvation mortality at *E*/*S* > 0.2 (consistent with Huss et al’s herring model) |
| *z_p_* | Parameter setting the decline in predation mortality as length increases | 8 |  |
| **Recruitment** | | | |
| $\alpha$ | Density-independent Ricker recruitment parameter | 1.2$\times$10^-1^ (herring-like, for use with spawning stock biomass tonnes), 8.4$\times$10^-9^ (cod-like, for use with total number eggs) | Lynam et al. (2005) for herring young-of-year recruitment data, Sparholt (1995) for cod |
| $\beta$ | Density-dependent Ricker recruitment parameter | 8.5$\times$10^-7^ (herring-like, for use with spawning stock biomass tonnes), 4.4$\times$10^-15^ (cod-like, for use with total number eggs) | See above |
| *p* | Egg packing density for conversion from mass to eggs (eggs/gram eggs) | 3.3$\times$10^6^ (for herring), 4.5$\times$10^6^ (cod-like) |  |
| $\sigma_{devo}$ | **Non-heritable noise in trait** | 3 levels (st. dev. 0 of trunc. Normal, st. dev. 1, st. dev. 3) | No non-heritable developmental stochasticity in trait values vs. low amount vs. high amount |
| **Resource base (see Equation 2)** | | | |
| *r_max_* | Logistic population growth rate of food resource | 1.5 |  |
| *K_half_* | Half-saturation constant for the Type II functional response of predators to the food resource | 1/10^th^ of initial *K_R_* |  |
| $\phi$ | Autocorrelation of *K_half_* between days in the model run | 0.8 |  |
| **Temperature dependence (see Equation 8)** | | | |
| *A* | Controls direction and amount of temperature dependence | 0.62 for *c_S_* and *c_E_*; 0.69 for $\rho_{1}$; -0.8 for *K_R_*; 0.73 for *r_max_* | Lindmark et al. (2022) |
| *T_ref_* | Baseline reference temperature | 284 Kelvin | Lindmark et al. (2022) |
| *k* | Boltzmann’s constant | 8.617$\times$10^-5^ | Lindmark et al. (2022) |

*References other than Audzijonyte and Richards (2018) or Huss et al (2012)

**Supplemental Table 2.** *Model parameters, descriptions of parameters, and values that were used together with sources of those values.*
