## Supplementary material for "Environmental variability alters and obscures fisheries-induced evolution": Code and simulated data files: extracting_output.R

```
object<-mclapply(c(1:98000), function(x) { model_iteration(x) }, mc.cores = 80 )

extract_output <- function(object, no_runs) {
data_list <- object # Use whatever the name is of the workspace object to which the results of the mclapply run were assigned 
no_runs <-no_runs # How many runs are contained in the object to which the mclapply results were assigned?
mat = matrix(, nrow = no_runs, ncol = 16) # Set up a matrix to hold the results
mapply(function(x) mapply(function(y) try(mat[x,y] <<- data_list[[x]][[y]]), y=c(1:6,12:13)), x=c(1:no_runs))
output_all <- as.data.frame(mat)
output_all <- output_all[,c(1:6,12:13)]
output_all <- na.omit(output_all)
colnames(output_all) <- c("phys", "AAM", "LAM", "L2", "L3", "Linf", "K_var", "F")
return(output_all)
}
```
