## Supplementary material for "Environmental variability alters and obscures fisheries-induced evolution": Code and simulated data files: plotting_code.R

```
# This performs the same job as the "extracting_output.R" file in the github repo
# The blocks of code after this block are written to make the figures from the "data" (simulation result) files which are already included in the data folder
data_list <- # object to which the mclapply results were assigned
no_runs <-21000
mat = matrix(, nrow = no_runs, ncol = 16)
mapply(function(x) mapply(function(y) try(mat[x,y] <<- data_list[[x]][[y]]), y=c(1:6,12:13)), x=c(1:no_runs))
output_all <- as.data.frame(mat)
output_all <- output_all[,c(1:6,12:13)]
colnames(output_all) <- c("phys", "AAM", "LAM", "L2", "L3", "Linf", "K_var", "F")
save_output_all <- rbind(save_output_all,output_all)
nrow(save_output_all)


# read in "cod_2trt_data.csv" and "herring_2trt_data.csv"
library(ggplot2)
library(viridis)
library(gridExtra)
library(cowplot)
library(metR)

# Fig 3
data_to_use <- cod_2trt_data
data_to_use <- data_to_use[which(data_to_use$phys!="NaN"),]
fish_tile <- aggregate(as.numeric(data_to_use$phys), by=list(as.numeric(data_to_use$K_var), as.numeric(data_to_use$F)), mean)
fish_tile[,2] <- log(exp(-1*fish_tile[,2])^(365))*-1
fish_tile[,1] <- sqrt(exp(fish_tile[,1]^2)-1)
fish_tile[,3] <- fish_tile[,3]/20
colnames(fish_tile) <- c("K_CV", "Fishing", "phys")
p1<-ggplot(fish_tile, aes(K_CV, Fishing, fill = phys, z=phys)) + geom_tile() + theme_minimal() + labs(x="Food resource variability", y="Fishing mort.", fill="Relative reproduction alloc.\nat short length") + scale_fill_viridis(option="turbo") + geom_contour(color="black", breaks=seq(.54,.67, by=.01)) + geom_text_contour(breaks=seq(.54,.67, by=.01)) + theme(legend.position = "none")
data_to_use <- herring_2trt_data
data_to_use <- data_to_use[which(data_to_use$phys!="NaN"),]
fish_tile <- aggregate(as.numeric(data_to_use$phys), by=list(as.numeric(data_to_use$K_var), as.numeric(data_to_use$F)), mean)
fish_tile[,2] <- log(exp(-1*fish_tile[,2])^(365))*-1
fish_tile[,1] <- sqrt(exp(fish_tile[,1]^2)-1)
fish_tile[,3] <- fish_tile[,3]/20
colnames(fish_tile) <- c("K_CV", "Fishing", "phys")
p2<-ggplot(fish_tile, aes(K_CV, Fishing, fill = phys, z=phys)) + geom_tile() + theme_minimal() + labs(x="Food resource variability", y="Fishing mort.", fill="Relative reproduction alloc.\nat short length") + scale_fill_viridis(option="turbo") + geom_contour(color="black", breaks=seq(.52,.70, by=.02)) + geom_text_contour(breaks=seq(.52,.70, by=.02)) + theme(legend.position = "none")
combined_plot <- plot_grid(p1,p2)
p2<-ggplot(fish_tile, aes(K_CV, Fishing, fill = phys, z=phys)) + geom_tile() + theme_minimal() + labs(x="Food resource variability", y="Fishing mort.", fill="Relative reproduction alloc.\nat short length") + scale_fill_viridis(option="turbo") + geom_contour(color="black", breaks=seq(.52,.70, by=.02)) + geom_text_contour(breaks=seq(.52,.70, by=.02))
legend <- get_legend(
  p2
)
plot_grid(combined_plot, legend)

# Combine combined plot and legend using plot_grid()
plot_grid(combined_plot, legend,ncol=1,rel_heights = c(1, .1))
p2<-ggplot(fish_tile, aes(K_CV, Fishing, fill = phys, z=phys)) + geom_tile() + theme_minimal() + labs(x="Food resource variability", y="Fishing mort.", fill="Relative reproduction alloc.\nat short length") + scale_fill_viridis(option="turbo") + geom_contour(color="black")


# Fig 5
data_to_use <- cod_2trt_data
data_to_use <- data_to_use[which(data_to_use$phys!="NaN"),]
fish_tile <- aggregate(as.numeric(data_to_use$LAM), by=list(as.numeric(data_to_use$K_var), as.numeric(data_to_use$F)), mean) # toggle the df name
fish_tile[,2] <- log(exp(-1*fish_tile[,2])^(365))*-1
fish_tile[,1] <- sqrt(exp(fish_tile[,1]^2)-1)
fish_tile[,3] <- fish_tile[,3]*100
colnames(fish_tile) <- c("K_CV", "Fishing", "phys")
p1<-ggplot(fish_tile, aes(K_CV, Fishing, fill = phys, z=phys)) + geom_tile() + theme_minimal() + labs(x="Food resource variability", y="Fishing mort.", fill="Length\nat maturity (cm.)") + scale_fill_viridis(option="turbo") + geom_contour(color="black") + geom_text_contour()
data_to_use <- herring_2trt_data
data_to_use <- data_to_use[which(data_to_use$phys!="NaN"),]
fish_tile <- aggregate(as.numeric(data_to_use$LAM), by=list(as.numeric(data_to_use$K_var), as.numeric(data_to_use$F)), mean) # toggle the df name
fish_tile[,2] <- log(exp(-1*fish_tile[,2])^(365))*-1
fish_tile[,1] <- sqrt(exp(fish_tile[,1]^2)-1)
fish_tile[,3] <- fish_tile[,3]*100
colnames(fish_tile) <- c("K_CV", "Fishing", "phys")
p2<-ggplot(fish_tile, aes(K_CV, Fishing, fill = phys, z=phys)) + geom_tile() + theme_minimal() + labs(x="Food resource variability", y="Fishing mort.", fill="Length\nat maturity (cm.)") + scale_fill_viridis(option="turbo") + geom_contour(color="black") + geom_text_contour()
grid.arrange(p1,p2, widths=c(1,1))


# Fig 4
data_to_use <- Fig4_table_cod
fish_tile <- aggregate(as.numeric(data_to_use$phys), by=list(data_to_use$base_K, as.numeric(data_to_use$F)), mean)
colnames(fish_tile) <- c("base_K", "Fishing", "phys")
fish_tile$Fishing <- as.numeric(fish_tile$Fishing)
fish_tile$phys <- as.numeric(fish_tile$phys)
fish_tile[,2] <- log(exp(-1*fish_tile[,2])^(365))*-1
std_mean <- function(x) sd(x)/sqrt(length(x))
fish_tile_se <- aggregate(as.numeric(data_to_use$phys), by=list(data_to_use$base_K, as.numeric(data_to_use$F)), std_mean)
colnames(fish_tile_se) <- c("base_K", "Fishing", "phys_se")
fish_tile_se$Fishing <- as.numeric(fish_tile_se$Fishing)
fish_tile_se$phys_se <- as.numeric(fish_tile_se$phys_se)
fish_tile[,4] <- fish_tile_se$phys_se
colnames(fish_tile)[4] <- "phys_se"
p1<-ggplot(fish_tile, aes(x=Fishing, y=phys/20, colour=base_K)) + theme_classic() +
  geom_line() +
  geom_point(aes(colour=base_K, shape=base_K)) + labs(y="Relative reproduction allocation\nat short length", x="Fishing mortality", col="Condition") + ylim(0.5,0.71) + theme(legend.position = "none")
data_to_use <- Fig4_table_herring 
fish_tile <- aggregate(as.numeric(data_to_use$phys), by=list(data_to_use$base_K, as.numeric(data_to_use$F)), mean)
colnames(fish_tile) <- c("base_K", "Fishing", "phys")
fish_tile$Fishing <- as.numeric(fish_tile$Fishing)
fish_tile$phys <- as.numeric(fish_tile$phys)
fish_tile[,2] <- log(exp(-1*fish_tile[,2])^(365))*-1
std_mean <- function(x) sd(x)/sqrt(length(x))
fish_tile_se <- aggregate(as.numeric(data_to_use$phys), by=list(data_to_use$base_K, as.numeric(data_to_use$F)), std_mean)
colnames(fish_tile_se) <- c("base_K", "Fishing", "phys_se")
fish_tile_se$Fishing <- as.numeric(fish_tile_se$Fishing)
fish_tile_se$phys_se <- as.numeric(fish_tile_se$phys_se)
fish_tile[,4] <- fish_tile_se$phys_se
colnames(fish_tile)[4] <- "phys_se"
p2 <- ggplot(fish_tile, aes(x=Fishing, y=phys/20, colour=base_K)) + theme_classic() +
  geom_line() +
  geom_point(aes(colour=base_K, shape=base_K)) + labs(y="", x="Fishing mortality", col="Condition", shape="Condition") + ylim(0.5,0.71) + theme(legend.position = "none")
combined_plot <- plot_grid(p1,p2)
p2 <- ggplot(fish_tile, aes(x=Fishing, y=phys/20, colour=base_K)) + theme_classic() +
  geom_line() +
  geom_point(aes(colour=base_K, shape=base_K)) + labs(y="", x="Fishing mortality", col="Condition", shape="Condition") + geom_errorbar(aes(ymin=phys/20-phys_se/20, ymax=phys/20+phys_se/20)) + ylim(0.5,0.71)
legend <- get_legend(
  p2
)
plot_grid(combined_plot, legend)

# Supplemental Figure 3 (yields with fishing)
yield <- read.csv("~/Documents/yield_SuppFig2.csv")
p1<-ggplot(yield[which(yield$species=="cod-like"),], aes(x=F_max, y=yield, colour=as.character(resource_var))) + theme_classic() +
geom_line() +
geom_point(aes(colour=as.character(resource_var), shape=as.character(resource_var))) + labs(y="Yield in biomass", x="Maximum fishing\nmortality", shape="Resource variability", col="Resource variability", title="Cod-like") + theme(legend.position = "none")
p2<-ggplot(yield[which(yield$species=="herring-like"),], aes(x=F_max, y=yield, colour=as.character(resource_var))) + theme_classic() +
geom_line() +
geom_point(aes(colour=as.character(resource_var), shape=as.character(resource_var))) + labs(y="Yield in biomass", x="Maximum fishing\nmortality", shape="Resource variability", col="Resource variability", title="Herring-like") + theme(legend.position = "none")
combined_plot <- plot_grid(p1,p2)
p2<-ggplot(yield[which(yield$species=="herring-like"),], aes(x=F_max, y=yield, colour=as.character(resource_var))) + theme_classic() +
  geom_line() +
  geom_point(aes(colour=as.character(resource_var), shape=as.character(resource_var))) + labs(y="Yield in biomass", x="Maximum fishing\nmortality", shape="Resource variability", col="Resource variability", title="Herring-like")
legend <- get_legend(
  p2
)
plot_grid(combined_plot, legend)

# Supplemental Figure 4 (yield/biomass, low vs high base K)
yield <- read.csv("~/Documents/yield_SuppFig3.csv")
yield$yield.per.bmass <- yield$yield.per.bmass*100
p1<-ggplot(yield[which(yield$species=="cod-like"),], aes(x=F_max, y=yield.per.bmass, colour=as.character(resource_var))) + theme_classic() +
  geom_line() +
  geom_point(aes(colour=as.character(resource_var), shape=as.character(resource_var))) + labs(y="Yield as % of popn. biomass", x="Maximum fishing\nmortality", shape="Resource variability", col="Resource variability", title="Cod-like") + theme(legend.position = "none")
p2<-ggplot(yield[which(yield$species=="herring-like"),], aes(x=F_max, y=yield.per.bmass, colour=as.character(resource_var))) + theme_classic() +
  geom_line() +
  geom_point(aes(colour=as.character(resource_var), shape=as.character(resource_var))) + labs(y="Yield as % of popn. biomass", x="Maximum fishing\nmortality", shape="Resource variability", col="Resource variability", title="Herring-like") + theme(legend.position = "none")
combined_plot <- plot_grid(p1,p2)
p2<-ggplot(yield[which(yield$species=="herring-like"),], aes(x=F_max, y=yield.per.bmass, colour=as.character(resource_var))) + theme_classic() +
  geom_line() +
  geom_point(aes(colour=as.character(resource_var), shape=as.character(resource_var))) + labs(y="Yield as % of popn. biomass", x="Maximum fishing\nmortality", shape="Resource variability", col="Resource variability", title="Herring-like")
legend <- get_legend(
  p2
)
plot_grid(combined_plot, legend, rel_widths = c(1,1,0.3))
```
