## Supplementary material for "Environmental variability alters and obscures fisheries-induced evolution": Code and simulated data files: plotting_code_thermal_analyses.R

```
# To hoover up .csv files generated by the modeling code and merge together into single .csv files with all data from a run
myMergedData <- 
  do.call(rbind,
          lapply(paste("data_storage/", list.files(paste(getwd(),"/data_storage", sep="") ), sep=""), read.csv))

library(ggplot2)
library(cowplot)
library(viridis)
library(metR)

##### Main text - Figure 6

data_to_use <- cod_posT
data_to_use <- data_to_use[which(data_to_use$phys!="NaN"),]
fish_tile <- aggregate(as.numeric(data_to_use$LAM), by=list(as.numeric(data_to_use$K_var), as.numeric(data_to_use$F)), mean) # toggle the df name
fish_tile[,2] <- log(exp(-1*fish_tile[,2])^(365))*-1
fish_tile[,1] <- sqrt(exp(fish_tile[,1]^2)-1)
fish_tile[,3] <- fish_tile[,3]*100
colnames(fish_tile) <- c("K_CV", "Fishing", "phys")
p_cod_posT<-ggplot(fish_tile, aes(K_CV, Fishing, fill = phys, z=phys)) + geom_tile()  + theme_minimal() + labs(x="Temperature variability, \nCV", y="Maximum fishing mort.", fill="Length\nat maturity (cm.)") + scale_fill_viridis(option="turbo") + geom_contour(color="black", lwd=0.2) + geom_text_contour(stroke=0.2) + scale_fill_viridis_c(limits=c(40,48), option = "turbo") + theme(legend.position = "none")

data_to_use <- cod_negT
data_to_use <- data_to_use[which(data_to_use$phys!="NaN"),]
fish_tile <- aggregate(as.numeric(data_to_use$LAM), by=list(as.numeric(data_to_use$K_var), as.numeric(data_to_use$F)), mean) # toggle the df name
fish_tile[,2] <- log(exp(-1*fish_tile[,2])^(365))*-1
fish_tile[,1] <- sqrt(exp(fish_tile[,1]^2)-1)
fish_tile[,3] <- fish_tile[,3]*100
colnames(fish_tile) <- c("K_CV", "Fishing", "phys")
p_cod_negT<-ggplot(fish_tile, aes(K_CV, Fishing, fill = phys, z=phys)) + geom_tile()  + theme_minimal() + labs(x="Temperature variability", y="Maximum fishing mort.", fill="Length\nat maturity (cm.)") + scale_fill_viridis(option="turbo") + geom_contour(color="black") + geom_text_contour() + scale_fill_viridis_c(limits=c(40,48), option = "turbo")
cod_legend<-get_legend(p_cod_negT)
data_to_use <- cod_negT
data_to_use <- data_to_use[which(data_to_use$phys!="NaN"),]
fish_tile <- aggregate(as.numeric(data_to_use$LAM), by=list(as.numeric(data_to_use$K_var), as.numeric(data_to_use$F)), mean) # toggle the df name
fish_tile[,2] <- log(exp(-1*fish_tile[,2])^(365))*-1
fish_tile[,1] <- sqrt(exp(fish_tile[,1]^2)-1)
fish_tile[,3] <- fish_tile[,3]*100
colnames(fish_tile) <- c("K_CV", "Fishing", "phys")
p_cod_negT<-ggplot(fish_tile, aes(K_CV, Fishing, fill = phys, z=phys)) + geom_tile()  + theme_minimal() + labs(x="Temperature variability, \nCV", y="Maximum fishing mort.", fill="Length\nat maturity (cm.)") + scale_fill_viridis(option="turbo") + geom_contour(color="black", lwd=0.2) + geom_text_contour(stroke=0.2) + scale_fill_viridis_c(limits=c(40,48), option = "turbo") + theme(legend.position = "none")

data_to_use <- herring_posT
data_to_use <- data_to_use[which(data_to_use$phys!="NaN"),]
fish_tile <- aggregate(as.numeric(data_to_use$LAM), by=list(as.numeric(data_to_use$K_var), as.numeric(data_to_use$F)), mean) # toggle the df name
fish_tile[,2] <- log(exp(-1*fish_tile[,2])^(365))*-1
fish_tile[,1] <- sqrt(exp(fish_tile[,1]^2)-1)
fish_tile[,3] <- fish_tile[,3]*100
colnames(fish_tile) <- c("K_CV", "Fishing", "phys")
p_herring_posT<-ggplot(fish_tile, aes(K_CV, Fishing, fill = phys, z=phys)) + geom_tile()  + theme_minimal() + labs(x="Temperature variability, \nCV", y="Maximum fishing mort.", fill="Length\nat maturity (cm.)") + scale_fill_viridis(option="turbo") + geom_contour(color="black", lwd=0.2, breaks=seq(27.1, 28.3, by=0.1)) + geom_text_contour(stroke=0.2, breaks=seq(27.1, 28.3, by=0.1)) + scale_fill_viridis_c(limits=c(26.9,28.4), option = "turbo") + theme(legend.position = "none")

data_to_use <- herring_negT
data_to_use <- data_to_use[which(data_to_use$phys!="NaN"),]
fish_tile <- aggregate(as.numeric(data_to_use$LAM), by=list(as.numeric(data_to_use$K_var), as.numeric(data_to_use$F)), mean) # toggle the df name
fish_tile[,2] <- log(exp(-1*fish_tile[,2])^(365))*-1
fish_tile[,1] <- sqrt(exp(fish_tile[,1]^2)-1)
fish_tile[,3] <- fish_tile[,3]*100
colnames(fish_tile) <- c("K_CV", "Fishing", "phys")
p_herring_negT<-ggplot(fish_tile, aes(K_CV, Fishing, fill = phys, z=phys)) + geom_tile()  + theme_minimal() + labs(x="Temperature variability", y="Maximum fishing mort.", fill="Length\nat maturity (cm.)") + scale_fill_viridis(option="turbo") + geom_contour(color="black") + geom_text_contour() + scale_fill_viridis_c(limits=c(26.9,28.4), option = "turbo") 
herring_legend<-get_legend(p_herring_negT)
data_to_use <- herring_negT
data_to_use <- data_to_use[which(data_to_use$phys!="NaN"),]
fish_tile <- aggregate(as.numeric(data_to_use$LAM), by=list(as.numeric(data_to_use$K_var), as.numeric(data_to_use$F)), mean) # toggle the df name
fish_tile[,2] <- log(exp(-1*fish_tile[,2])^(365))*-1
fish_tile[,1] <- sqrt(exp(fish_tile[,1]^2)-1)
fish_tile[,3] <- fish_tile[,3]*100
colnames(fish_tile) <- c("K_CV", "Fishing", "phys")
p_herring_negT<-ggplot(fish_tile, aes(K_CV, Fishing, fill = phys, z=phys)) + geom_tile()  + theme_minimal() + labs(x="Temperature variability, \nCV", y="Maximum fishing mort.", fill="Length\nat maturity (cm.)") + scale_fill_viridis(option="turbo") + geom_contour(color="black", lwd=0.2, breaks=seq(26.9, 27.5, by=0.1)) + geom_text_contour(stroke=0.2, breaks=seq(26.9, 27.5, by=0.1)) + scale_fill_viridis_c(limits=c(26.9,28.4), option = "turbo") + theme(legend.position = "none")

plot_grid(p_cod_posT, p_cod_negT, cod_legend, p_herring_posT, p_herring_negT, herring_legend, nrow=2, rel_widths = c(1,1,0.1,1,1,0.1))

##### Supplemental Fig 5

# length at 3 years of age

data_to_use <- cod_posT
data_to_use <- data_to_use[which(data_to_use$phys!="NaN"),]
fish_tile <- aggregate(as.numeric(data_to_use$L3), by=list(as.numeric(data_to_use$K_var), as.numeric(data_to_use$F)), mean) # toggle the df name
fish_tile[,2] <- log(exp(-1*fish_tile[,2])^(365))*-1
fish_tile[,1] <- sqrt(exp(fish_tile[,1]^2)-1)
fish_tile[,3] <- fish_tile[,3]*100
colnames(fish_tile) <- c("K_CV", "Fishing", "phys")
p_cod_posT_L3<-ggplot(fish_tile, aes(K_CV, Fishing, fill = phys, z=phys)) + geom_tile()  + theme_minimal() + labs(x="Temperature variability, \nCV", y="Maximum fishing mort.", fill="Length\nat 3 yrs. (cm.)") + scale_fill_viridis(limits=c(46,51),option="turbo") + geom_contour(color="black", lwd=0.2) + geom_text_contour(stroke=0.2) + scale_fill_viridis_c( limits=c(46,51),option = "turbo") + theme(legend.position = "none")

data_to_use <- cod_negT
data_to_use <- data_to_use[which(data_to_use$phys!="NaN"),]
fish_tile <- aggregate(as.numeric(data_to_use$L3), by=list(as.numeric(data_to_use$K_var), as.numeric(data_to_use$F)), mean) # toggle the df name
fish_tile[,2] <- log(exp(-1*fish_tile[,2])^(365))*-1
fish_tile[,1] <- sqrt(exp(fish_tile[,1]^2)-1)
fish_tile[,3] <- fish_tile[,3]*100
colnames(fish_tile) <- c("K_CV", "Fishing", "phys")
p_cod_negT<-ggplot(fish_tile, aes(K_CV, Fishing, fill = phys, z=phys)) + geom_tile()  + theme_minimal() + labs(x="Temperature variability", y="Maximum fishing mort.", fill="Length\nat 3 yrs. (cm.)") + scale_fill_viridis(limits=c(46,51),option="turbo") + geom_contour(color="black") + geom_text_contour() + scale_fill_viridis_c(option = "turbo")
cod_legend_L3<-get_legend(p_cod_negT)
data_to_use <- cod_negT
data_to_use <- data_to_use[which(data_to_use$phys!="NaN"),]
fish_tile <- aggregate(as.numeric(data_to_use$L3), by=list(as.numeric(data_to_use$K_var), as.numeric(data_to_use$F)), mean) # toggle the df name
fish_tile[,2] <- log(exp(-1*fish_tile[,2])^(365))*-1
fish_tile[,1] <- sqrt(exp(fish_tile[,1]^2)-1)
fish_tile[,3] <- fish_tile[,3]*100
colnames(fish_tile) <- c("K_CV", "Fishing", "phys")
p_cod_negT_L3<-ggplot(fish_tile, aes(K_CV, Fishing, fill = phys, z=phys)) + geom_tile()  + theme_minimal() + labs(x="Temperature variability, \nCV", y="Maximum fishing mort.", fill="Length\nat 3 yrs. (cm.)") + scale_fill_viridis(option="turbo") + geom_contour(color="black", lwd=0.2, breaks=seq(46.4, 48.2, by=0.2)) + geom_text_contour(stroke=0.2, breaks=seq(46.4, 48.2, by=0.2)) + scale_fill_viridis_c(limits=c(46,51), option = "turbo") + theme(legend.position = "none")


data_to_use <- herring_posT
data_to_use <- data_to_use[which(data_to_use$phys!="NaN"),]
fish_tile <- aggregate(as.numeric(data_to_use$L3), by=list(as.numeric(data_to_use$K_var), as.numeric(data_to_use$F)), mean) # toggle the df name
fish_tile[,2] <- log(exp(-1*fish_tile[,2])^(365))*-1
fish_tile[,1] <- sqrt(exp(fish_tile[,1]^2)-1)
fish_tile[,3] <- fish_tile[,3]*100
colnames(fish_tile) <- c("K_CV", "Fishing", "phys")
p_herring_posT_L3<-ggplot(fish_tile, aes(K_CV, Fishing, fill = phys, z=phys)) + geom_tile()  + theme_minimal() + labs(x="Temperature variability, \nCV", y="Maximum fishing mort.", fill="Length\nat 3 yrs. (cm.)") + scale_fill_viridis(limits=c(28,32),option="turbo") + geom_contour(color="black", lwd=0.2, breaks=seq(29,31.8, by=0.4)) + geom_text_contour(stroke=0.2, breaks=seq(29,31.8, by=0.4)) + scale_fill_viridis_c(limits=c(28,32), option = "turbo") + theme(legend.position = "none")

data_to_use <- herring_negT
data_to_use <- data_to_use[which(data_to_use$phys!="NaN"),]
fish_tile <- aggregate(as.numeric(data_to_use$L3), by=list(as.numeric(data_to_use$K_var), as.numeric(data_to_use$F)), mean) # toggle the df name
fish_tile[,2] <- log(exp(-1*fish_tile[,2])^(365))*-1
fish_tile[,1] <- sqrt(exp(fish_tile[,1]^2)-1)
fish_tile[,3] <- fish_tile[,3]*100
colnames(fish_tile) <- c("K_CV", "Fishing", "phys")
p_herring_negT<-ggplot(fish_tile, aes(K_CV, Fishing, fill = phys, z=phys)) + geom_tile()  + theme_minimal() + labs(x="Temperature variability", y="Maximum fishing mort.", fill="Length\nat 3 yrs. (cm.)") + scale_fill_viridis(option="turbo") + geom_contour(color="black") + geom_text_contour() + scale_fill_viridis_c(limits=c(28,32), option = "turbo") 
herring_legend_L3<-get_legend(p_herring_negT)
data_to_use <- herring_negT
data_to_use <- data_to_use[which(data_to_use$phys!="NaN"),]
fish_tile <- aggregate(as.numeric(data_to_use$L3), by=list(as.numeric(data_to_use$K_var), as.numeric(data_to_use$F)), mean) # toggle the df name
fish_tile[,2] <- log(exp(-1*fish_tile[,2])^(365))*-1
fish_tile[,1] <- sqrt(exp(fish_tile[,1]^2)-1)
fish_tile[,3] <- fish_tile[,3]*100
colnames(fish_tile) <- c("K_CV", "Fishing", "phys")
p_herring_negT_L3<-ggplot(fish_tile, aes(K_CV, Fishing, fill = phys, z=phys)) + geom_tile()  + theme_minimal() + labs(x="Temperature variability, \nCV", y="Maximum fishing mort.", fill="Length\nat 3 yrs. (cm.)") + scale_fill_viridis(option="turbo") + geom_contour(color="black", lwd=0.2, breaks=seq(29,30.6, by=0.2)) + geom_text_contour(stroke=0.2, breaks=seq(29,30.6, by=0.2)) + scale_fill_viridis_c(limits=c(28,32), option = "turbo") + theme(legend.position = "none")

# age at maturation

data_to_use <- cod_posT
data_to_use <- data_to_use[which(data_to_use$phys!="NaN"),]
fish_tile <- aggregate(as.numeric(data_to_use$AAM), by=list(as.numeric(data_to_use$K_var), as.numeric(data_to_use$F)), mean) # toggle the df name
fish_tile[,2] <- log(exp(-1*fish_tile[,2])^(365))*-1
fish_tile[,1] <- sqrt(exp(fish_tile[,1]^2)-1)
fish_tile[,3] <- (fish_tile[,3]-1)/4
colnames(fish_tile) <- c("K_CV", "Fishing", "phys")
p_cod_posT_age<-ggplot(fish_tile, aes(K_CV, Fishing, fill = phys, z=phys)) + geom_tile()  + theme_minimal() + labs(x="Temperature variability, \nCV", y="Maximum fishing mort.", fill="Age\nat maturity (years)") + scale_fill_viridis(option="turbo") + geom_contour(color="black", lwd=0.2, breaks=seq(2.25, 2.9, by=0.05)) + geom_text_contour(stroke=0.2, breaks=c(2.25, 2.9, by=0.05)) + scale_fill_viridis_c(limits=c(2.2,3), option = "turbo") + theme(legend.position = "none")

data_to_use <- cod_negT
data_to_use <- data_to_use[which(data_to_use$phys!="NaN"),]
fish_tile <- aggregate(as.numeric(data_to_use$AAM), by=list(as.numeric(data_to_use$K_var), as.numeric(data_to_use$F)), mean) # toggle the df name
fish_tile[,2] <- log(exp(-1*fish_tile[,2])^(365))*-1
fish_tile[,1] <- sqrt(exp(fish_tile[,1]^2)-1)
fish_tile[,3] <- (fish_tile[,3]-1)/4
colnames(fish_tile) <- c("K_CV", "Fishing", "phys")
p_cod_negT<-ggplot(fish_tile, aes(K_CV, Fishing, fill = phys, z=phys)) + geom_tile()  + theme_minimal() + labs(x="Temperature variability", y="Maximum fishing mort.", fill="Age\nat maturity (years)") + scale_fill_viridis(limits=c(2.2,3),option="turbo") + geom_contour(color="black") + geom_text_contour() + scale_fill_viridis_c( option = "turbo")
cod_legend_age<-get_legend(p_cod_negT)
data_to_use <- cod_negT
data_to_use <- data_to_use[which(data_to_use$phys!="NaN"),]
fish_tile <- aggregate(as.numeric(data_to_use$AAM), by=list(as.numeric(data_to_use$K_var), as.numeric(data_to_use$F)), mean) # toggle the df name
fish_tile[,2] <- log(exp(-1*fish_tile[,2])^(365))*-1
fish_tile[,1] <- sqrt(exp(fish_tile[,1]^2)-1)
fish_tile[,3] <- (fish_tile[,3]-1)/4
colnames(fish_tile) <- c("K_CV", "Fishing", "phys")
p_cod_negT_age<-ggplot(fish_tile, aes(K_CV, Fishing, fill = phys, z=phys)) + geom_tile()  + theme_minimal() + labs(x="Temperature variability, \nCV", y="Maximum fishing mort.", fill="Age\nat maturity (years)") + scale_fill_viridis(limits=c(2.2,3),option="turbo") + geom_contour(color="black", lwd=0.2, breaks=seq(2.25, 2.9, by=0.05)) + geom_text_contour(stroke=0.2, breaks=c(2.25, 2.9, by=0.05)) + scale_fill_viridis_c( option = "turbo") + theme(legend.position = "none")


data_to_use <- herring_posT
data_to_use <- data_to_use[which(data_to_use$phys!="NaN"),]
fish_tile <- aggregate(as.numeric(data_to_use$AAM), by=list(as.numeric(data_to_use$K_var), as.numeric(data_to_use$F)), mean) # toggle the df name
fish_tile[,2] <- log(exp(-1*fish_tile[,2])^(365))*-1
fish_tile[,1] <- sqrt(exp(fish_tile[,1]^2)-1)
fish_tile[,3] <- (fish_tile[,3]-1)/4
colnames(fish_tile) <- c("K_CV", "Fishing", "phys")
p_herring_posT_age<-ggplot(fish_tile, aes(K_CV, Fishing, fill = phys, z=phys)) + geom_tile()  + theme_minimal() + labs(x="Temperature variability, \nCV", y="Maximum fishing mort.", fill="Age\nat maturity (years)") + scale_fill_viridis(option="turbo") + geom_contour(color="black", lwd=0.2) + geom_text_contour(stroke=0.2) + scale_fill_viridis_c( option = "turbo") + theme(legend.position = "none")

data_to_use <- herring_negT
data_to_use <- data_to_use[which(data_to_use$phys!="NaN"),]
fish_tile <- aggregate(as.numeric(data_to_use$AAM), by=list(as.numeric(data_to_use$K_var), as.numeric(data_to_use$F)), mean) # toggle the df name
fish_tile[,2] <- log(exp(-1*fish_tile[,2])^(365))*-1
fish_tile[,1] <- sqrt(exp(fish_tile[,1]^2)-1)
fish_tile[,3] <- (fish_tile[,3]-1)/4
colnames(fish_tile) <- c("K_CV", "Fishing", "phys")
p_herring_negT<-ggplot(fish_tile, aes(K_CV, Fishing, fill = phys, z=phys)) + geom_tile()  + theme_minimal() + labs(x="Temperature variability", y="Maximum fishing mort.", fill="Age\nat maturity (years)") + scale_fill_viridis(option="turbo") + geom_contour(color="black") + geom_text_contour() + scale_fill_viridis_c(option = "turbo") 
herring_legend_age<-get_legend(p_herring_negT)
data_to_use <- herring_negT
data_to_use <- data_to_use[which(data_to_use$phys!="NaN"),]
fish_tile <- aggregate(as.numeric(data_to_use$AAM), by=list(as.numeric(data_to_use$K_var), as.numeric(data_to_use$F)), mean) # toggle the df name
fish_tile[,2] <- log(exp(-1*fish_tile[,2])^(365))*-1
fish_tile[,1] <- sqrt(exp(fish_tile[,1]^2)-1)
fish_tile[,3] <- (fish_tile[,3]-1)/4
colnames(fish_tile) <- c("K_CV", "Fishing", "phys")
p_herring_negT_age<-ggplot(fish_tile, aes(K_CV, Fishing, fill = phys, z=phys)) + geom_tile()  + theme_minimal() + labs(x="Temperature variability, \nCV", y="Maximum fishing mort.", fill="Age\nat maturity (years)") + scale_fill_viridis(option="turbo") + geom_contour(color="black", lwd=0.2) + geom_text_contour(stroke=0.2) + scale_fill_viridis_c( option = "turbo") + theme(legend.position = "none")

# Evolution of the genetically controlled parameter r

data_to_use <- cod_posT
data_to_use <- data_to_use[which(data_to_use$phys!="NaN"),]
fish_tile <- aggregate(as.numeric(data_to_use$phys), by=list(as.numeric(data_to_use$K_var), as.numeric(data_to_use$F)), mean)
fish_tile[,2] <- log(exp(-1*fish_tile[,2])^(365))*-1
fish_tile[,1] <- sqrt(exp(fish_tile[,1]^2)-1)
fish_tile[,3] <- fish_tile[,3]/20
colnames(fish_tile) <- c("K_CV", "Fishing", "phys")
p_cod_posT_evo<-ggplot(fish_tile, aes(K_CV, Fishing, fill = phys, z=phys)) + geom_tile()  + theme_minimal() + labs(x="Temperature variability, \nCV", y="Maximum fishing mort.", fill="r, relative reproduction alloc.\nat short length") + scale_fill_viridis(limits=c(0.53,0.68),option="turbo") + geom_contour(color="black", lwd=0.2, breaks=seq(0.54,0.67, by=0.01)) + geom_text_contour(stroke=0.2, breaks=seq(0.54,0.67, by=0.01)) + scale_fill_viridis_c( limits=c(0.53,0.68),option = "turbo") + theme(legend.position = "none")

data_to_use <- cod_negT
data_to_use <- data_to_use[which(data_to_use$phys!="NaN"),]
fish_tile <- aggregate(as.numeric(data_to_use$phys), by=list(as.numeric(data_to_use$K_var), as.numeric(data_to_use$F)), mean)
fish_tile[,2] <- log(exp(-1*fish_tile[,2])^(365))*-1
fish_tile[,1] <- sqrt(exp(fish_tile[,1]^2)-1)
fish_tile[,3] <- fish_tile[,3]/20
colnames(fish_tile) <- c("K_CV", "Fishing", "phys")
p_cod_negT<-ggplot(fish_tile, aes(K_CV, Fishing, fill = phys, z=phys)) + geom_tile()  + theme_minimal() + labs(x="Temperature variability", y="Maximum fishing mort.", fill="r, relative reproduction alloc.\nat short length") + scale_fill_viridis(limits=c(0.53,0.68),option="turbo") + geom_contour(color="black") + geom_text_contour() + scale_fill_viridis_c(limits=c(0.53,0.68), option = "turbo")
cod_legend_evo<-get_legend(p_cod_negT)
data_to_use <- cod_negT
data_to_use <- data_to_use[which(data_to_use$phys!="NaN"),]
fish_tile <- aggregate(as.numeric(data_to_use$phys), by=list(as.numeric(data_to_use$K_var), as.numeric(data_to_use$F)), mean)
fish_tile[,2] <- log(exp(-1*fish_tile[,2])^(365))*-1
fish_tile[,1] <- sqrt(exp(fish_tile[,1]^2)-1)
fish_tile[,3] <- fish_tile[,3]/20
colnames(fish_tile) <- c("K_CV", "Fishing", "phys")
p_cod_negT_evo<-ggplot(fish_tile, aes(K_CV, Fishing, fill = phys, z=phys)) + geom_tile()  + theme_minimal() + labs(x="Temperature variability, \nCV", y="Maximum fishing mort.", fill="r, relative reproduction alloc.\nat short length") + scale_fill_viridis(limits=c(0.53,0.68),option="turbo") + geom_contour(color="black", lwd=0.2, breaks=seq(0.54,0.67, by=0.01)) + geom_text_contour(stroke=0.2, breaks=seq(0.54,0.67, by=0.01)) + scale_fill_viridis_c(limits=c(0.53,0.68), option = "turbo") + theme(legend.position = "none")


data_to_use <- herring_posT
data_to_use <- data_to_use[which(data_to_use$phys!="NaN"),]
fish_tile <- aggregate(as.numeric(data_to_use$phys), by=list(as.numeric(data_to_use$K_var), as.numeric(data_to_use$F)), mean)
fish_tile[,2] <- log(exp(-1*fish_tile[,2])^(365))*-1
fish_tile[,1] <- sqrt(exp(fish_tile[,1]^2)-1)
fish_tile[,3] <- fish_tile[,3]/20
colnames(fish_tile) <- c("K_CV", "Fishing", "phys")
p_herring_posT_evo<-ggplot(fish_tile, aes(K_CV, Fishing, fill = phys, z=phys)) + geom_tile()  + theme_minimal() + labs(x="Temperature variability, \nCV", y="Maximum fishing mort.", fill="r, relative reproduction alloc.\nat short length") + scale_fill_viridis(limits=c(0.5, 0.75),option="turbo") + geom_contour(color="black", lwd=0.2, breaks=seq(0.52,0.68,by=0.02)) + geom_text_contour(stroke=0.2, breaks=seq(0.52,0.68,by=0.02)) + scale_fill_viridis_c(limits=c(0.5, 0.75),  option = "turbo") + theme(legend.position = "none")

data_to_use <- herring_negT
data_to_use <- data_to_use[which(data_to_use$phys!="NaN"),]
fish_tile <- aggregate(as.numeric(data_to_use$phys), by=list(as.numeric(data_to_use$K_var), as.numeric(data_to_use$F)), mean)
fish_tile[,2] <- log(exp(-1*fish_tile[,2])^(365))*-1
fish_tile[,1] <- sqrt(exp(fish_tile[,1]^2)-1)
fish_tile[,3] <- fish_tile[,3]/20
colnames(fish_tile) <- c("K_CV", "Fishing", "phys")
p_herring_negT<-ggplot(fish_tile, aes(K_CV, Fishing, fill = phys, z=phys)) + geom_tile()  + theme_minimal() + labs(x="Temperature variability", y="Maximum fishing mort.", fill="r, relative reproduction alloc.\nat short length") + scale_fill_viridis(limits=c(0.5, 0.75),option="turbo") + geom_contour(color="black") + geom_text_contour() + scale_fill_viridis_c( limits=c(0.5, 0.75), option = "turbo") 
herring_legend_evo<-get_legend(p_herring_negT)
data_to_use <- herring_negT
data_to_use <- data_to_use[which(data_to_use$phys!="NaN"),]
fish_tile <- aggregate(as.numeric(data_to_use$phys), by=list(as.numeric(data_to_use$K_var), as.numeric(data_to_use$F)), mean)
fish_tile[,2] <- log(exp(-1*fish_tile[,2])^(365))*-1
fish_tile[,1] <- sqrt(exp(fish_tile[,1]^2)-1)
fish_tile[,3] <- fish_tile[,3]/20
colnames(fish_tile) <- c("K_CV", "Fishing", "phys")
p_herring_negT_evo<-ggplot(fish_tile, aes(K_CV, Fishing, fill = phys, z=phys)) + geom_tile()  + theme_minimal() + labs(x="Temperature variability, \nCV", y="Maximum fishing mort.", fill="r, relative reproduction alloc.\nat short length") + scale_fill_viridis(limits=c(0.5, 0.75), option="turbo") + geom_contour(color="black", lwd=0.2, breaks=seq(0.52,0.7,by=0.02)) + geom_text_contour(stroke=0.2, breaks=seq(0.52,0.7,by=0.02)) + scale_fill_viridis_c( limits=c(0.5, 0.75),option = "turbo") + theme(legend.position = "none")

thermal_plot_L3<-plot_grid(p_cod_posT_L3, p_cod_negT_L3, cod_legend_L3, p_herring_posT_L3, p_herring_negT_L3, herring_legend_L3, nrow=1, rel_width=c(1,1,0.1, 1,1, 0.1))

thermal_plot_age<-plot_grid(p_cod_posT_age, p_cod_negT_age, cod_legend_age, p_herring_posT_age, p_herring_negT_age, herring_legend_age, nrow=1, rel_width=c(1,1,0.1, 1,1, 0.1))

thermal_plot_evo<-plot_grid(p_cod_posT_evo, p_cod_negT_evo, cod_legend_evo, p_herring_posT_evo, p_herring_negT_evo, herring_legend_evo, nrow=1, rel_width=c(1,1,0.1, 1,1, 0.1))
```
