## Supplementary material for "Environmental variability alters and obscures fisheries-induced evolution": Code and simulated data files: growth_functions_backup.R

```
# GROWTH FUNCTIONS

# For obtaining structural mass
structural_mass <- function(l) {
  return(c_1*(l)^c_2)
}

# Calculate the intrinsic length-dependent E/S ratio that the fish is trying to attain
intrinsic_lambda <- function(l, l_bar, r, lambda_min, lambda_max) {
  return(lambda_min + (lambda_max-lambda_min)*(exp(r*(l-l_bar)))/(1+exp(r*(l-l_bar))))
}

# Energy intake for the fish per year, dependent on fish structural biomass
size_dependent_energy_intake <- function(S, p_0, p_1) {
  return(p_0*(S)^(p_1))
}

# Calculate the cost of the year of maintaining current structural and reversible biomass
maintenance_cost <- function(S, E, c_S, c_E) {
  return(c_S*S + c_E*E)
}

# Function for calculating a new, reduced level of reversible biomass, if net energy intake is negative
E_for_maintenance <- function(E, p_net, e_E) {
  E[p_net < 0] <- E[p_net < 0] + p_net[p_net < 0]*(1/e_E)
  E[E<0] <- 0
  return(E)
}

# Calculate the amount of energy needed to raise the E/S ratio to the length-dependent maximum
p_E_raising_lambda <- function(lambda_l, S, E) {
  return((lambda_l*S - E))
}

# Calculating proportion of any remaining energy to be converted to reversible mass, to maintain E/S
remaining_energy_allocation_E <- function(e_S, e_E, lambda_l) {
  return((lambda_l*e_S)/(lambda_l*e_S + e_E))
}

# Same as above, but for structural mass
remaining_energy_allocation_S <- function(e_S, e_E, lambda_l) {
  return(1-(lambda_l*e_S)/(lambda_l*e_S + e_E))
}

get_pre_reproductive_size <- function(E_past, l_past, resource, new_mu_exp, consumed_resource, nu) {
  # Get the current structural mass of a cohort-phenotype combination
  S_past = structural_mass(l_past)
  # Create the values of structural mass S and reversible mass E to be updated over the course of the simulation
  S = S_past
  E = E_past
  # Calculate what the optimal ration of E/S (which the fish will attempt to optimize) is, given the length and the physiological phenotype
  lambda_l = intrinsic_lambda(l_past, l_bar, r, lambda_min, lambda_max)
  consumed_resource <- consumed_resource
  # Calculating energy intake and subtracting maintenance costs
  # Dividing the total resources consumed during the feeding bout (happens each day before growth and mortality, using the ode function in the model_run() function in "model_run_code") by nu, the sum of all fishes' maximum energy intakes, to determine how much this fish ate (it will be less than maximum energy intake because of competition and the Holling Type II consumption)
  p_net = size_dependent_energy_intake(S_past, p_0, p_1)*(consumed_resource/nu) - maintenance_cost(S_past, E_past, c_S, c_E)
  # Subtracting any negative net energy intakes from reversible masses for those cohort-pheno combos
  E = E_for_maintenance(E, p_net, e_E)
  # Setting the negative net energy intakes to 0 now that the costs have been paid
  p_net[p_net < 0] <- 0
  # Calculate the amount of energy that must be added to E for E/S to reach the optimum
  p_E <- p_E_raising_lambda(lambda_l, S, E)
  # Calculate by how much the net energy intake exceeds the amount needed to bring lambda to the optimum
  surplus <- (p_E*(1/e_E)) - p_net
  surplus <- surplus*(-1)
  # In cases where the net energy intake does NOT exceed the amount needed to bring lambda to the optimum, all the remaining energy intake is added to E
  E[surplus <= 0] <- p_net[surplus <= 0]*e_E + E[surplus <= 0]
  # ... and, for those cohort-pheno combos, the amount of energy intake left over for them this day is set to zero
  p_net[surplus <= 0] <- 0
  # In cases where the net energy intake DOES exceed the amount needed to bring lambda to the optimum, p_E is added to E, and taken out of the remaining energy intake
  E[surplus > 0] <- p_E[surplus > 0] + E[surplus > 0]
  p_net[surplus > 0] <- p_net[surplus > 0] - p_E[surplus > 0]*(1/e_E)
  # Any remaining energy intake is spread between E and S in the way that does not change E/S
  E[surplus > 0] = E[surplus > 0] + e_E*p_net[surplus > 0]*remaining_energy_allocation_E(e_S, e_E, lambda_l)[surplus > 0]
  S[surplus > 0] = S[surplus > 0] + e_S*p_net[surplus > 0]*remaining_energy_allocation_S(e_S, e_E, lambda_l)[surplus > 0]
  l = (S*(1/c_1))^(1/c_2)
  output <- list(E,l,consumed_resource)
  return(output)
}
```
