## Supplementary material for "Environmental variability alters and obscures fisheries-induced evolution": Code and simulated data files: herring_same_structure_reproduction_functions_ML.R

```
# REPRODUCTION FUNCTIONS

fecundity <- function(E, l, mu_exp) {
  # Calculate potential reproductive contribution in terms of biomass, with all of the indirect costs of spawning (movement, behavioral, etc.) subtracted
  repro_biomass<-(w*E-r_0*structural_mass(l)^(r_1))*mu_exp
  # Next calculate the total amount that the fish would be spending on spawning, including both direct costs of egg and indirect costs
  repro_biomass_individual<-w*E
  # Zero out cohorts that have a negative amount of energy for eggs after the energy for indirect costs is taken away. These fish don't spawn
  repro_biomass_individual[repro_biomass < 0] <- 0 
  repro_biomass[repro_biomass < 0] <- 0 
  E <- E - repro_biomass_individual
  # Calculate spawning stock biomass in grams
  total_fecundity = sum(E[repro_biomass > 0]*mu_exp[repro_biomass > 0]) + sum(structural_mass(l[repro_biomass > 0])*mu_exp[repro_biomass > 0]) 
  # in tonnes
  total_fecundity = total_fecundity*(1e-6)
  return(list(total_fecundity, repro_biomass, E))
}

biomass_t <- function(E, l, mu_exp) {
  masses <- E + structural_mass(l)
  biomass <- sum(mu_exp*masses)
  return(biomass)
}

Ricker <- function(egg_production_total) {
  # Ricker for YOY
  N_0 = Alpha * egg_production_total * exp(-b*egg_production_total)
  # Converting from millions recruits to recruits, dividing by 2 because Ricker makes both male and female fish
  N_0 = N_0 * 1000000
  N_0 = N_0/2
  return(N_0)
}

inheritance <- function(repro_biomass_cohorts, mu, mu_exp, N_0) {
  # The 2D matrix (of fecundities of each cohort-phenotype combo; cohorts are rows, columns are phenotype) is expanded into an 3D array where each z-level is just a duplicate of the 2D matrix of fecundities, in order to then element-wise-multiply each cell by the corresponding proportions in the mu 3D array (in which each z-level is a locus, and a cell is the proportion of individuals in that cohort-pheno combo that have the '1' allele for that locus)	
  # The end result of the multiplication is a 3D array where each z level contains the expected number of gametes created by each cohort-phenotype combo (of those gametes that WILL be represented among the successful new recruits) that have that z-level's (that locus's) '1' allele
  multiplier<-array(c(rep(repro_biomass_cohorts,ncol(repro_biomass_cohorts))), dim=c(dim(repro_biomass_cohorts),ncol(repro_biomass_cohorts)))
  # Weighting the expected number of gametes from each cohort-pheno combo by the abundance of parents in that cohort-pheno combo has ALREADY happened, in the first line of code of the fecundity function
  allele1_contributions <- mu*multiplier
  # Produces a vector of total expected allele-1 carrying gametes from each locus, by summing on each z-level of the 3D array (i.e., each locus)  
  allele1_contributions_vec <- apply(allele1_contributions, 3, sum)
  # Produces a vector of total gametes produced (whether allele is 0 or 1), for each locus (essentially, a vector of the same value repeated multiple times, where the number of times is the number of loci). The previous vector generated can be divided by this to get expected proportion allele-1 gametes for each vector
  total_contributions_vec <- apply(multiplier, 3, sum)
  allele1_proportions <- allele1_contributions_vec/total_contributions_vec
  # The next few lines collapse the new recruits into "superindividuals" to speed computation time. If the number of new recruits N_0 is 1000 or larger, it will be divided by 10^(floor(log10(N_0)) - 2) (for N_0 of 1,000-9,999, this is 10, for N_0 of 10,000-99,999, this is 100, and so on). This will allow us to assign genotypes more quickly. The total number of recruits will be resized to its original order of magnitude later
  divider <- 1
  if (floor(log10(N_0)) > 2) {
    divider <- 10^(floor(log10(N_0)) - 2)
  }
  # This line uses expected allele-1 gametes for each locus to (for each locus) generate (using the Binomial distribution) actual gametes for that locus in the new recruits pool. This creates a matrix where columns are loci, rows are "super"-individuals, and matrix cells/entries are alleles	
  # Each "super"-individual's genotype is actually its expected gametic contribution to the next generation, as opposed to explicitly simulating diploid individuals
  new_genotypes<-mapply(function(x) rbinom(N_0/divider,1,x), allele1_proportions)
  # From the previous line, allele-0 gametes are represented as zeroes and allele-1 gametes as 1. This line will add across loci in order to, for each superindividual in the new recruit population, create a value between 0 and 1 representing the number of allele-1 locus each new recruit (or "super" recruit) has across all of its simulated loci. This will then be used to create a physiological reaction norm phenotype (but the phenotype is not a perfect function of this genotype, because developmental noise exists)
  new_genotypes_vec <- apply(new_genotypes, 1, sum)
  # The rtruncnorm function is used to (within the range of possible phenotypes) create a phenotype from the genotype with a little added noise. The 'round' function will discretize these phenotypes
  expression_list <- mapply(function(z) round(rtruncnorm(1,1,ncol(repro_biomass_cohorts),mean=z,sd=1)), new_genotypes_vec)
  expression_list <- as.vector(unlist(expression_list))
  # This line calculates the numbers of "super" recruits per phenotype using the tabulate function, then re-multiplies by the "divider" value to go back to units of total recruits
  expression_vec<-tabulate(expression_list, nbins=loci)*divider
  # This line will make the vector of phenotype frequencies (generated in the previous line) the first row (youngest cohort) of the new "mu expressed" matrix   
  mu_exp[1,] <- expression_vec
  return(list(mu, expression_list, new_genotypes, mu_exp))
}

scroll_through_phenotypes <- function(expressed, genotype) {
  # Create a row for each z-level in the new mu 3D array, where the row has the proportions of new recruits from each phenotype (column) that have the 1 allele for that locus  
  row<-mapply(function(y) length(expressed[which(expressed==y & genotype==1)])/length(expressed[which(expressed==y)]), seq(1,loci))
  row[is.na(row)] <- 0
  return(row)
}
```
