## Supplementary material for "Environmental variability alters and obscures fisheries-induced evolution": Code and simulated data files: model_run_code_herring_same_structure_ML.R

```
### HERRING-LIKE MODEL

# Dependent packages
library(pbapply)
library(truncnorm)
library(parallel)
library(deSolve)

# Setting up the crosses between different levels of the harvest and resource variability variables
rand_resource_vec <- rep((sqrt(log(seq(0,8, length=14)^2 + 1))), each=7000,times=1) # Values input to seq() are in units of coefficient of variation of lognormal distribution of resource K - they are back-converted here to SD of a Normal
fishing_vec <- rep(seq(0,1.8,by=1.8/13),each=500, times=14)

# The function that performs a given run of the model
model_iteration <<- function(results_counter) {
  # Dependent packages
  library(pbapply)
  library(truncnorm)
  
  # Run files containing functions for growth, reproduction and mortality processes
  
  source("mortality_functions.R")
  
  source("herring_same_structure_reproduction_functions_ML.R")
  
  source("growth_functions_backup.R")
  
  # Set longevity of the fish species, runtime of the model, and number of loci
  a_max_days <<- 5475
  # set runtime to any arbitrary desired value, in units of days, that is a multiple of 365
  runtime_days <<-  100*365 # 100*365 for 36500 days = 100 years
  loci <<- 20
  # Setting the length of the timestep in days - set to any fraction of 365. These are, by default, seasons of the year (365/4 days long)
  timescale <<-  (365)/4
  # Setting run time and longevity to units of time steps
  runtime <<- round(runtime_days/timescale)
  a_max <<- round(a_max_days/timescale)
  # Whether fish breeds once a year (has a breeding season), or breeds continously throughout the year except for some period of time
  # Cod and herring both have a breeding season, meaning they spawn in just one time step out of year
  breeding_season <<- TRUE
  # How long to run model for convergence before addition of genetic diversity
  # in units of days
  add_genotypes <<- 0.6*runtime_days
  
  # Set parameters
  # The steepness of the increase in the ratio lambda (prioritizing reversible over structural mass) as the fish grows longer
  r <<-  6
  # The length at which the increasing function of lambda reaches its inflection point
  l_bar <<- 0.2
  # Scaling parameter for obtaining structural mass from standard length
  # From Huss et al
  c_1 <<- 5735
  c_2 <<-  3.125
  # Multiplier for obtaining overall daily energy intake as a function of structural mass
  # See Mitchell Jones' 2014 UVM thesis and Rudstam cited within - NOT considering increased costs (and increased consumption) during travel (which our model does explicitly model), this value is reasonable
  p_0 <<-  0.3
  # Exponent for obtaining overall daily energy intake as a function of structural mass
  p_1 <<-  2/3
  # Maximum ratio lambda (reversible over structural mass) a fish can attain over its life
  # From Huss et al
  lambda_max <<- 1.13
  # Minimum ratio lambda a fish can attain over its life
  lambda_min <<-  0
  # Multiplier for obtaining cost of maintenance of structural mass
  # From Huss et al
  c_S <<-  0.03 
  # Multiplier for obtaining cost of maintenance of reversible mass
  c_E <<-  0.03 
  # Efficiency with which energy is converted into reversible mass
  e_E <<-  0.9
  # Efficiency with which energy is converted to structural mass
  e_S <<- (1/3)
  # Multiplier and exponent for the cost of reproduction (in grams)
  # Megrey et al Ecological Modelling 2007 and other sources put 15-25% total mass lost as typical for reproducing herring, and Huss et al uses 0.5 conversion efficiency of reproductive costs to gametes
  # Wheeler 2009 says that 30cm is typical length for herring at 50% probability of being mature, which would be S=133.2102 and E=93.24714 (for K=0.7), and mass for all reproduction costs would be 46.6, but halve that for the 0.5 efficiency from Huss
  r_0 <<- 1.25
  r_1 <<- 0.6
  # Effectively, reproductive investment (fraction of reversible mass devoted to reproduction)
  # The 20% fraction of *total* mass lost during reproduction, and K=0.7 for maturity (Megrey, Rutsdam, Slotte 1999), gives us the fraction of *reversible* mass E specifically that will be spent on reproduction
  w<<- 0.5
  # Instantaneous base daily mortality rate, derived from daily instantaneous probability of survival for M of 0.2. Multiplied by -1 to make positive.
  m <<- log(0.999452)*(-1)
  # Instantaneous fishing mortality
  Fishing <<- 0
  # For what fishing level to add when fishing is introduced partway through the simulation - this expression takes yearly instantaneous fishing mortality, gives probability of surviving the year
  Fishing_add <<- exp(-1*fishing_vec[results_counter])
  Fishing_add <<- log((Fishing_add)^(1/365))*(-1)
  # Ricker
  # Lynam MEPS 2005
  Alpha <<- 1.16e-1
  b <<- 8.53e-7
  # Parameter setting size-selection ogive steepness
  q_f <<- 0.5
  # Minimum catch size for the species (in meters)
  mincatchsize <<- 0.3
  # Increased natural mortality rate at low E/S or low S  
  # For roughly 0.5 year-based instantaneous mortality at ~1 yrs. from small-size-based predation mortality, Köster et al 2003
  # https://images.app.goo.gl/HNzVWX17UJEpSiM68
  # exp(-0.5) = 0.6065307 = 60.65307% remaining after 1 yr.
  # 0.6065307^(1/365) = 0.9986
  m_p_max<<-log(0.9986)*(-1)
  # Maximum condition dependent mortality for a daily instantaneous survival probaility of 5% (or anything in 1-10% range) guarantees instantaneous probability of survival of <2% for any fish with truly zero body condition - this is the right order of magnitude for fish to have virtually no chance of survival if their reversible mass is nil
  m_c_max<<-log(0.95)*(-1)
  # ^ similar modeling papers generally need to assess a range of mortality values for sensitivity and exact ideal values to use are uncertain (e.g., Audzijonyte and Richards 2018 p E156)  
  # Egg packing constant
  packing <<- 3333333
  # Parameter for exponential decline in condition-dependent mortality with improved body condition
  # Huss et al - a ~5 times faster decline in starvation mortality with increased condition than for the cod-like model, so that this mortality is negligible when E/S > 0.2 (Huss et al)
  z_c<<-35
  # Parameter for exponential decline in predation mortality with increased body length
  z_p<<-8
  # Amount of resource (initial), set to reasonable starting values by calculating fish population resource consumption
  resource <<- 3e+14
  # Resource carrying capacity
  resource_K <<- 3e+14
  # Resource intrinsic growth rate
  resource_r <<- 1.5
  # Resource variability
  r_SD <<- rand_resource_vec[results_counter]
  # Consumer functional response half-saturation constant
  K_half <<- 3e+13
  
  # Initializing the matrices for the first time point
  timepoints <<- vector(mode = "list", length = runtime)
  # Each timepoint consists of a matrix known as E of reversible masses, organized in rows for ages and columns for physiological phenotypes (i.e., what is their function for lambda with length); a matrix of the same organization, named l, for the lengths of fish in each cohort-phenotype combination; a matrix of the same organization for numbers or densities of individuals, mu_exp (for "expressed"), where each column is a different phenotype (differences between the phenotypes before genotypic differences are added will be very minimal and only due to developmental noise); and an array where each z-level is a locus and the cells contain the proportion of each age-phenotype combo that has the '1' allele for that locus  
  timepoint1 <<- list(matrix(data=0, nrow=a_max, ncol=loci), 
                      matrix(data=0, nrow=a_max, ncol=loci),
                      matrix(data=0, nrow=a_max, ncol=loci),
                      array(data=0, dim=c(a_max, loci, loci)))
  # YOY (young of year) individuals sizes and lengths - the model begins with a set number of YOY individuals	
  # YOY initial reversible E and structural S biomas is treated as fixed and unchanging and unrelated to genotype affecting adult growth  
  # Arrhenius and Hansson ICES 1996, for herring-like
  timepoint1[[1]][1,] <<- 0.3712037
  timepoint1[[2]][1,] <<- 0.057
  # Setting initial number of individuals
  timepoint1[[3]][1,] <<- 100
  # Initially, every cohort is euqally likely to have a 1 or 0 at any locus  
  timepoint1[[4]][1,,] <<- 0.5
  timepoints[[1]] <<- timepoint1
  # Initialize resources vector
  resources <<- vector(mode="numeric", length=runtime)
  # Initialize resource consumption vector
  resource_consumption <<- vector(mode="numeric", length=runtime)
  # Initialize vector to hold yields
  yields <<- vector(mode="numeric", length=runtime)
  # This list will hold summary statistics of random values generated for the resource carrying capacity at every time period/season of the model run	
  random_vec_summaries <<- vector(mode="list", length=runtime)
  # Initialize vector of recruitments
  recruitments <<- vector(mode="numeric", length=runtime)
  
  # Iterate through time steps - this actually runs the model
  pbmapply( function(x) {
    timepoints[[x]] <<- model_run(timepoints[[x-1]], x)
  }, 2:runtime)
  
  # Initialiizing a matrix
  phenotype_averages <<- matrix(nrow=6, ncol=loci)
  
  for (tracker_counter in 1:loci) {
    # This matrix determines which cohort-pheno combos are mature. The model is stopped right before the last reproduction would occur, so we can use preparedness to mature (this is given by equation 7 of the paper methods) as an index of maturity 
    mature<<-(w*as.vector(timepoints[[runtime]][[1]][,tracker_counter])-as.vector((r_0*(structural_mass(timepoints[[runtime]][[2]][,tracker_counter]))^(r_1))))
    # This line determines what the earliest age is at which members of a phenotype are mature
    phenotype_averages[1,tracker_counter] <<- which(mature==(mature[mature>0])[1])
    # This line determines the length of the individuals of the youngest-to-be-mature cohort within each phenotype group
    phenotype_averages[2,tracker_counter] <<- timepoints[[runtime]][[2]][phenotype_averages[1,tracker_counter],tracker_counter]
    # This line determines the total number of individuals of the youngest-to-be-mature cohort within each phenotype group
    phenotype_averages[3,tracker_counter] <<- timepoints[[runtime]][[3]][phenotype_averages[1,tracker_counter],tracker_counter]
    # This line determines the length of age-2 individuals within each phenotype group
    phenotype_averages[4,tracker_counter] <<- timepoints[[runtime]][[2]][9,tracker_counter]
    # This line determines the length of age-3 individuals within each phenotype group
    phenotype_averages[5,tracker_counter] <<- timepoints[[runtime]][[2]][13,tracker_counter]
    # This line determines the length of age-10 individuals within each phenotype group
    phenotype_averages[6,tracker_counter] <<- timepoints[[runtime]][[2]][41,tracker_counter]
  }
  
  # Turning abundances of each newly-mature cohort within each phenotypic group into proportions of newly mature fish across phenotypic groups
  phenotype_averages[3,] <<- phenotype_averages[3,]/sum(phenotype_averages[3,])
  
  # Calculating average value of the physiological phenotype across the phenotypic groups, using the abundance proportions generated in the previous line.
  result_inflection <- sum(c(1:loci)*(timepoints[[runtime-(365/timescale)]][[3]][1,]/sum(timepoints[[runtime-(365/timescale)]][[3]][1,])))
  # Calculating average age at maturity
  result_AAM <- sum(phenotype_averages[3,]*phenotype_averages[1,])
  # Calculating average length at maturity
  result_LAM <- sum(phenotype_averages[3,]*phenotype_averages[2,])
  # Calculating average length at age 2
  result_L2 <- sum(phenotype_averages[3,]*phenotype_averages[4,])
  # Calculating average length at age 3
  result_L3 <- sum(phenotype_averages[3,]*phenotype_averages[5,])
  # Calculating average length at age 10 (probably not "L_inf", but close to the longest length the fish will achieve in its life)
  result_Linf <- sum(phenotype_averages[3,]*phenotype_averages[6,])
  
  # Returning values generated and time series
  result_popn <-mapply(function(x) sum(timepoints[[x]][[3]]), seq((365/timescale)*10,runtime, by=365/timescale))
  result_biomass <-mapply(function(x) sum((timepoints[[x]][[1]] + timepoints[[x]][[2]])*timepoints[[x]][[3]] ), seq(40,400, by=4))
  return(list(result_inflection, result_AAM, result_LAM, result_L2,result_L3, result_Linf, result_popn, result_biomass, resources, resource_consumption, yields, r_SD, Fishing_add, random_vec_summaries, recruitments, as.data.frame(phenotype_averages)))
}

# Function for an iteration of the model
model_run <-function(list, x) {
  # Adding fishing after a certain time point
  if (x==round(add_genotypes/timescale)) {
    Fishing <<- Fishing_add
  }
  # Adding differences between genotypes
  if (x==round(add_genotypes/timescale)) {
    r <<- seq(0, 12, by = (12-0)/(loci-1))
    r<<-matrix(rep(r,each=a_max),nrow=a_max)
  }
  # Creating the new E, l, and mu_exp matrices to be filled
  new_E_l <<- list(list[[1]], list[[2]])
  new_mu_exp <<- list[[3]]
  # Generate vectors of random values, with some degree of temporal correlation phi
  times <- 1:round(timescale)
  mat <- as.matrix(dist(times, diag = T, upper= T))
  # Set the autocorrelation below
  phi = 0.8
  # Creating the correlation-by-time-separation matrix
  AR_mat <- as.matrix(phi^mat)
  L <- t(chol(AR_mat))
  # Multiplying the transpose of the Cholesky factorization by a vector of random values gives vectors of autocorrelated value
  ar_var <- L%*%matrix(rnorm(round(timescale)*round(timescale),0,1), ncol= round(timescale))*r_SD*sqrt(1-phi^2)
  random_values <- as.vector(ar_var[,1])
  random_vec_summaries[[x]] <<- summary(exp(random_values - (sd(random_values)^2)/2))
  mapply ( function(growing) {
    # Total maximum energy consumptions across all fish of different sizes
    nu<-sum(size_dependent_energy_intake(structural_mass(new_E_l[[2]]), p_0, p_1)*new_mu_exp)
    # Compute a random value for the resource's carrying capacity
    random_value <- random_values[growing]
    random_resource_K <- resource_K*(exp(random_value - (sd(random_values)^2)/2))
    # Allow the fish to feed (with their maximum food intake dependent on their sizes the previous day) and the resource to simultaneously grow
    # Differential equation function
    Feeding <- function(t, state, parameters) {
      with(as.list(c(state, parameters)), {
        dR <- resource_r*R*((resource_K-R)/resource_K) - nu*((R)/(R+K_half))
        dC <- nu*((R)/(R+K_half))
        list(c(dR, dC))
      })
    }
    # Parameters for the DE
    parameters <- c(resource_r = resource_r, resource_K = random_resource_K, K_half=K_half, nu=nu)
    # States of resource density and resource consumed
    state <- c(R=resource, C=0)
    times <- seq(0, 1, by = 0.1)
    # Numerically integrate the differential equations
    out <- ode(y = state, times = times, func = Feeding, parms = parameters)  
    if (as.data.frame(out)$R[nrow(out)]<2) {
      print(out)
    }
    resource <<- as.data.frame(out)$R[nrow(out)]
    # Here, the fish grow
    new_E_l <<- get_pre_reproductive_size(new_E_l[[1]], new_E_l[[2]], resource, new_mu_exp, as.data.frame(out)$C[nrow(out)], nu)
    # New population sizes for each cohort-genotype combo, after mortality and prior to breeding
    before_fishing <- biomass_t(new_E_l[[1]], new_E_l[[2]], new_mu_exp)
    temp_new_mu_exp <<- fishing_mortality(new_mu_exp, new_E_l[[1]], new_E_l[[2]])
    after_fishing <- biomass_t(new_E_l[[1]], new_E_l[[2]], temp_new_mu_exp)
    yield <<- before_fishing - after_fishing
    new_mu_exp <<- mortality(new_mu_exp, new_E_l[[1]], new_E_l[[2]])
  }, 1:round(timescale))
  # Record resource level
  resources[x] <<- resource
  # Record consumption level
  # Resource consumption is only for last day within period of days in 'timescale' (i.e., the last day of the season if, timescale = 365/4)  
  resource_consumption[x] <<- new_E_l[[3]]
  # Record yield
  yields[x] <<- yield
  # Add oldest and second-oldest cohorts to make the 'plus group'	  
  new_mu_exp[nrow(new_mu_exp)-1,] <- new_mu_exp[nrow(new_mu_exp),] + new_mu_exp[nrow(new_mu_exp)-1,]
  # Empty out first row for young of year, for the mu_exp, E and l matrices
  new_mu_exp <- advance_age(new_mu_exp)
  new_E_l[[1]] <- advance_age(new_E_l[[1]])
  new_E_l[[2]] <- advance_age(new_E_l[[2]])
  # Empty out first row for young of year, for the mu matrix
  mu <- list[[4]]
  mu[2:nrow(mu),,] <- mu[1:(nrow(mu)-1),,]
  mu[1,,] <- 0
  # Reproduction is seasonal
  if ((breeding_season==T) & x%%round(365/timescale ) == 0 & x!=runtime) {
    # Get the amount of eggs produced and the amounts produced by each cohort-genotype combo, and a new E after reversible energy is depleted by mating
    fecundity_object <- fecundity(new_E_l[[1]], new_E_l[[2]], new_mu_exp)
    # The updated E matrix with cohorts' individuals' reversible mass values depleted (if they mated)
    new_E_l[[1]] <- fecundity_object[[3]]
    # Update first row of mu_exp with new individuals with expression based on past genotype
    new_recruits <- Ricker(fecundity_object[[1]])
    # Record the total number of new recruits
    recruitments[x] <<- new_recruits
    if (new_recruits>0) {
    inheritance_object <- inheritance(fecundity_object[[2]], mu, new_mu_exp, Ricker(fecundity_object[[1]]))
    # Updating mu_exp matrix
    new_mu_exp <- inheritance_object[[4]]
    # Give the YOY their sizes and lengths, if there are any
    if (sum(new_mu_exp[1,]) > 0) {
      new_E_l[[1]][1,] <- 0.3712037
      new_E_l[[2]][1,] <- 0.057
    }
    # Create vectors with length=number of new individuals, where each value is the individual's expressed phenotype (for "expressed") and their genotype (for "genotype")
    # "genotype" has one of these for each locus (for each locus, giving the alleles that each of the individuals have for that locus)
    mu <- inheritance_object[[1]]
    expressed <- inheritance_object[[2]]
    genotype <- inheritance_object[[3]]
    # length(expressed) should == length(genotype)
    # Filling in the genotypes of the new recruits
    # The scroll_through_genotypes function will create a row for each z-level in the new mu 3D array, where the row has the proportions of new recruits from each phenotype (column) that have the '1' allele for the locus that this z-level of the array corresponds to    
    # See "reproduction_function" file for more info in comments
    new_cohort_genotypes<-mapply(function(k) scroll_through_phenotypes(expressed, genotype[,k]), seq(1, loci))    
    mu[1,,] <- new_cohort_genotypes
  }
  }
  # Return updated matrices for this time point (2 through runtime)
  return(list(new_E_l[[1]], new_E_l[[2]], new_mu_exp, mu))
}

# Function for advancing age
advance_age <- function(matrix) {
  matrix[2:(nrow(matrix)),] <- matrix[1:(nrow(matrix)-1),]
  matrix[1,] <- 0
  return(matrix)
}
```
