## Supplementary material for "Environmental variability alters and obscures fisheries-induced evolution": Code and simulated data files: model_run_code_herring_same_structure_ML_thermal.R

```
### HERRING-LIKE MODEL - thermal stochasticity

# Dependent packages
library(pbapply)
library(truncnorm)
library(parallel)
library(deSolve)

# NOTE! some of the references in comments to "resources" may may actually pertain to temperature because they are unchanged from the version of this code which does NOT use thermal variation, as this code file uses the same code to vary temperature stochastically rather than directly varying food resources.
# Setting up the crosses between different levels of the harvest and temperature variability variables
rand_resource_vec <- rep((sqrt(log(seq(0,2, length=12)^2 + 1))), each=4900,times=1) # Values input to seq() are in units of coefficient of variation of lognormal distribution of temperatures - they are back-converted here to SD of a Normal
fishing_vec <- rep(seq(0,1.8,by=1.8/13),each=350, times=12)

# Arrhenius function for temperature dependencies in rates
Arr <- function(temp, A_v) {
  # the hard-coded value is Boltzmann's constant
  # temperatures in Kelvin
  exp((A_v*(temp-Tref))/((8.617e-5)*(temp*Tref)))
}

# The function that performs a given run of the model
model_iteration <<- function(results_counter) {
  # Dependent packages
  library(pbapply)
  library(truncnorm)
  
  # Run files containing functions for growth, reproduction and mortality processes
  
  source("mortality_functions.R")
  
  source("herring_same_structure_reproduction_functions_ML.R")
  
  source("growth_functions_backup.R")
  
  # Set longevity of the fish species, runtime of the model, and number of loci
  a_max_days <<- 5475
  # set runtime to any arbitrary desired value, in units of days, that is a multiple of 365
  runtime_days <<-  100*365 # 100*365 for 36500 days = 100 years
  loci <<- 20
  # Setting the length of the timestep in days - set to any fraction of 365. These are, by default, seasons of the year (365/4 days long)
  timescale <<-  (365)/4
  # Setting run time and longevity to units of time steps
  runtime <<- round(runtime_days/timescale)
  a_max <<- round(a_max_days/timescale)
  # Whether fish breeds once a year (has a breeding season), or breeds continuously throughout the year except for some period of time
  # Cod and herring both have a breeding season, meaning they spawn in just one time step out of year
  breeding_season <<- TRUE
  # How long to run model for convergence before addition of genetic diversity
  # in units of days
  add_genotypes <<- 0.6*runtime_days
  
  # Set parameters
  # The steepness of the increase in the ratio lambda (prioritizing reversible over structural mass) as the fish grows longer
  r <<-  6
  # The length at which the increasing function of lambda reaches its inflection point
  l_bar <<- 0.2
  # Scaling parameter for obtaining structural mass from standard length
  # From Huss et al
  c_1 <<- 5735
  c_2 <<-  3.125
  # Multiplier for obtaining overall daily energy intake as a function of structural mass
  # See Mitchell Jones' 2014 UVM thesis and Rudstam cited within - NOT considering increased costs (and increased consumption) during travel (which our model does explicitly model), this value is reasonable
  p_0 <<-  0.3
  # Exponent for obtaining overall daily energy intake as a function of structural mass
  p_1 <<-  2/3
  # Maximum ratio lambda (reversible over structural mass) a fish can attain over its life
  # From Huss et al
  lambda_max <<- 1.13
  # Minimum ratio lambda a fish can attain over its life
  lambda_min <<-  0
  # Multiplier for obtaining cost of maintenance of structural mass
  # From Huss et al
  c_S <<-  0.03 
  # Multiplier for obtaining cost of maintenance of reversible mass
  c_E <<-  0.03 
  # Efficiency with which energy is converted into reversible mass
  e_E <<-  0.9
  # Efficiency with which energy is converted to structural mass
  e_S <<- (1/3)
  # Multiplier and exponent for the cost of reproduction (in grams)
  # Megrey et al Ecological Modelling 2007 and other sources put 15-25% total mass lost as typical for reproducing herring, and Huss et al uses 0.5 conversion efficiency of reproductive costs to gametes
  # Wheeler 2009 says that 30cm is typical length for herring at 50% probability of being mature, which would be S=133.2102 and E=93.24714 (for K=0.7), and mass for all reproduction costs would be 46.6, but halve that for the 0.5 efficiency from Huss
  r_0 <<- 1.25
  r_1 <<- 0.6
  # Effectively, reproductive investment (fraction of reversible mass devoted to reproduction)
  # The 20% fraction of *total* mass lost during reproduction, and K=0.7 for maturity (Megrey, Rutsdam, Slotte 1999), gives us the fraction of *reversible* mass E specifically that will be spent on reproduction
  w<<- 0.5
  # Instantaneous base daily mortality rate, derived from daily instantaneous probability of survival for M of 0.2. Multiplied by -1 to make positive.
  m <<- log(0.999452)*(-1)
  # Instantaneous fishing mortality
  Fishing <<- 0
  # For what fishing level to add when fishing is introduced partway through the simulation - this expression takes yearly instantaneous fishing mortality, gives probability of surviving the year
  Fishing_add <<- exp(-1*fishing_vec[results_counter])
  Fishing_add <<- log((Fishing_add)^(1/365))*(-1)
  # Ricker
  # Lynam MEPS 2005
  Alpha <<- 1.16e-1
  b <<- 8.53e-7
  # Parameter setting size-selection ogive steepness
  q_f <<- 0.5
  # Minimum catch size for the species (in meters)
  mincatchsize <<- 0.3
  # Increased natural mortality rate at low E/S or low S  
  # For roughly 0.5 year-based instantaneous mortality at ~1 yrs. from small-size-based predation mortality, Köster et al 2003
  # https://images.app.goo.gl/HNzVWX17UJEpSiM68
  # exp(-0.5) = 0.6065307 = 60.65307% remaining after 1 yr.
  # 0.6065307^(1/365) = 0.9986
  m_p_max<<-log(0.9986)*(-1)
  # Maximum condition dependent mortality for a daily instantaneous survival probaility of 5% (or anything in 1-10% range) guarantees instantaneous probability of survival of <2% for any fish with truly zero body condition - this is the right order of magnitude for fish to have virtually no chance of survival if their reversible mass is nil
  m_c_max<<-log(0.95)*(-1)
  # ^ similar modeling papers generally need to assess a range of mortality values for sensitivity and exact ideal values to use are uncertain (e.g., Audzijonyte and Richards 2018 p E156)  
  # Egg packing constant
  packing <<- 3333333
  # Parameter for exponential decline in condition-dependent mortality with improved body condition
  # Huss et al - a ~5 times faster decline in starvation mortality with increased condition than for the cod-like model, so that this mortality is negligible when E/S > 0.2 (Huss et al)
  z_c<<-35
  # Parameter for exponential decline in predation mortality with increased body length
  z_p<<-8
  # Amount of resource (initial), set to reasonable starting values by calculating fish population resource consumption
  resource <<- 3e+14
  # Resource carrying capacity
  resource_K <<- 3e+14
  # Resource intrinsic growth rate
  resource_r <<- 1.5
  # Resource variability
  r_SD <<- rand_resource_vec[results_counter]
  # Consumer functional response half-saturation constant
  K_half <<- 3e+13
  
  # Environmental noise autocorrelation
  phi <<- 0.8
  
  # Reference Kelvin temperature for thermal modeling
  Tref <<- 284
  A_met <<- 0.62
  A_mor <<- 0.62
  A_int <<- 0.69
  A_gro <<- 0.73
  A_car <<- -0.8
  
  # Initializing the matrices for the first time point
  timepoints <<- vector(mode = "list", length = runtime)
  # Each timepoint consists of a matrix known as E of reversible masses, organized in rows for ages and columns for physiological phenotypes (i.e., what is their function for lambda with length); a matrix of the same organization, named l, for the lengths of fish in each cohort-phenotype combination; a matrix of the same organization for numbers or densities of individuals, mu_exp (for "expressed"), where each column is a different phenotype (differences between the phenotypes before genotypic differences are added will be very minimal and only due to developmental noise); and an array where each z-level is a locus and the cells contain the proportion of each age-phenotype combo that has the '1' allele for that locus  
  timepoint1 <<- list(matrix(data=0, nrow=a_max, ncol=loci), 
                      matrix(data=0, nrow=a_max, ncol=loci),
                      matrix(data=0, nrow=a_max, ncol=loci),
                      array(data=0, dim=c(a_max, loci, loci)))
  # YOY (young of year) individuals sizes and lengths - the model begins with a set number of YOY individuals	
  # YOY initial reversible E and structural S biomas is treated as fixed and unchanging and unrelated to genotype affecting adult growth  
  # Arrhenius and Hansson ICES 1996, for herring-like
  timepoint1[[1]][1,] <<- 0.3712037
  timepoint1[[2]][1,] <<- 0.057
  # Setting initial number of individuals
  timepoint1[[3]][1,] <<- 100
  # Initially, every cohort is euqally likely to have a 1 or 0 at any locus  
  timepoint1[[4]][1,,] <<- 0.5
  timepoints[[1]] <<- timepoint1
  # Initialize resources vector
  resources <<- vector(mode="numeric", length=runtime)
  # Initialize resource consumption vector
  resource_consumption <<- vector(mode="numeric", length=runtime)
  # Initialize vector to hold yields
  yields <<- vector(mode="numeric", length=runtime)
  # This list will hold summary statistics of random values generated for the resource carrying capacity at every time period/season of the model run	
  random_vec_summaries <<- vector(mode="list", length=runtime)
  # Initialize vector of recruitments
  recruitments <<- vector(mode="numeric", length=runtime)
  
  # Initialize holder for random values for environmental stochasticity
  ar_var<<-vector(mode = "list", length = round(runtime/12))
  
  # Iterate through time steps - this actually runs the model
  pbmapply( function(x) {
    timepoints[[x]] <<- model_run(timepoints[[x-1]], x)
  }, 2:runtime)
  
  # Initializing a matrix
  phenotype_averages <<- matrix(nrow=6, ncol=loci)
  
  for (tracker_counter in 1:loci) {
    # This matrix determines which cohort-pheno combos are mature. The model is stopped right before the last reproduction would occur, so we can use preparedness to mature (this is given by equation 7 of the paper methods) as an index of maturity 
    mature<<-(w*as.vector(timepoints[[runtime]][[1]][,tracker_counter])-as.vector((r_0*(structural_mass(timepoints[[runtime]][[2]][,tracker_counter]))^(r_1))))
    # This line determines what the earliest age is at which members of a phenotype are mature
    phenotype_averages[1,tracker_counter] <<- which(mature==(mature[mature>0])[1])
    # This line determines the length of the individuals of the youngest-to-be-mature cohort within each phenotype group
    phenotype_averages[2,tracker_counter] <<- timepoints[[runtime]][[2]][phenotype_averages[1,tracker_counter],tracker_counter]
    # This line determines the total number of individuals of the youngest-to-be-mature cohort within each phenotype group
    phenotype_averages[3,tracker_counter] <<- timepoints[[runtime]][[3]][phenotype_averages[1,tracker_counter],tracker_counter]
    # This line determines the length of age-2 individuals within each phenotype group
    phenotype_averages[4,tracker_counter] <<- timepoints[[runtime]][[2]][9,tracker_counter]
    # This line determines the length of age-3 individuals within each phenotype group
    phenotype_averages[5,tracker_counter] <<- timepoints[[runtime]][[2]][13,tracker_counter]
    # This line determines the length of age-10 individuals within each phenotype group
    phenotype_averages[6,tracker_counter] <<- timepoints[[runtime]][[2]][41,tracker_counter]
  }
  
  # Turning abundances of each newly-mature cohort within each phenotypic group into proportions of newly mature fish across phenotypic groups
  phenotype_averages[3,] <<- phenotype_averages[3,]/sum(phenotype_averages[3,])
  
  # Calculating average value of the physiological phenotype across the phenotypic groups, using the abundance proportions generated in the previous line.
  result_inflection <- sum(c(1:loci)*(timepoints[[runtime-(365/timescale)]][[3]][1,]/sum(timepoints[[runtime-(365/timescale)]][[3]][1,])))
  # Calculating average age at maturity
  result_AAM <- sum(phenotype_averages[3,]*phenotype_averages[1,])
  # Calculating average length at maturity
  result_LAM <- sum(phenotype_averages[3,]*phenotype_averages[2,])
  # Calculating average length at age 2
  result_L2 <- sum(phenotype_averages[3,]*phenotype_averages[4,])
  # Calculating average length at age 3
  result_L3 <- sum(phenotype_averages[3,]*phenotype_averages[5,])
  # Calculating average length at age 10 (probably not "L_inf", but close to the longest length the fish will achieve in its life)
  result_Linf <- sum(phenotype_averages[3,]*phenotype_averages[6,])
  
  # Returning values generated and time series
  result_popn <-mapply(function(x) sum(timepoints[[x]][[3]]), seq((365/timescale)*10,runtime, by=365/timescale))
  result_biomass <-mapply(function(x) sum((timepoints[[x]][[1]] + timepoints[[x]][[2]])*timepoints[[x]][[3]] ), seq(40,400, by=4))
  # After each single run, save output to a file
  # Get rid of the return() statement?
  object<-(list(result_inflection, result_AAM, result_LAM, result_L2, result_L3, result_Linf, result_popn, result_biomass, resources, resource_consumption, yields, r_SD, Fishing_add, random_vec_summaries, recruitments, as.data.frame(phenotype_averages)))
  
  data_list <- object # Use whatever the name is of the workspace object to which the results of the mclapply run were assigned
  no_runs <-1 # How many runs are contained in the object to which the mclapply results were assigned?
  mat = matrix(, nrow = no_runs, ncol = 16) # Set up a matrix to hold the results
  mapply(function(y) try(mat[1,y] <<- data_list[[y]]), y=c(1:6,12:13))
  output_all <- as.data.frame(mat)
  output_all <- output_all[,c(1:6,12:13)]
  output_all <- na.omit(output_all)
  colnames(output_all) <- c("phys", "AAM", "LAM", "L2", "L3", "Linf", "K_var", "F")
  
  # Unlike the versions of these code files which simulated variability in food resources, this code is designed to write output files to a folder as they are generated, rather than generating one large R object with all of the data to be analyzed by the "extracting_output.R" file. 
  # To run these files on a different machine/system/configuration, directories names' below must be changed.
  # Code for processing the .csv files generated by these is in the "plotting_code_thermal_analyses.R" file in the "figure_data+code" folder.
  
  setwd("/home/mschumm/ecoevoharvestmodel/data_storage")
  
  write.csv(output_all, paste("output_all_", results_counter, sep=""))
  
  setwd("/home/mschumm/ecoevoharvestmodel")
  
  return()
}

# Function for an iteration of the model
model_run <-function(list, x) {
  
  p_0 <<-  0.3
  c_S <<-  0.03
  c_E <<-  0.03
  m <<- log(0.999452)*(-1)
  resource_K <<- 3e+14
  resource_r <<- 1.5
  
  # Adding fishing after a certain time point
  if (x==round(add_genotypes/timescale)) {
    Fishing <<- Fishing_add
  }
  # Adding differences between genotypes
  if (x==round(add_genotypes/timescale)) {
    r <<- seq(0, 12, by = (12-0)/(loci-1))
    r<<-matrix(rep(r,each=a_max),nrow=a_max)
  }
  # Creating the new E, l, and mu_exp matrices to be filled
  new_E_l <<- list(list[[1]], list[[2]])
  new_mu_exp <<- list[[3]]
  if (((x-1)%%12 == 1) & x!=runtime) {
    # Generate vectors of random values, with some degree of temporal correlation phi
    times <- 1:(round(timescale)*12)
    mat <- as.matrix(dist(times, diag = T, upper= T))
    # Set the autocorrelation below
    # Creating the correlation-by-time-separation matrix
    AR_mat <- as.matrix(phi^mat)
    L <- t(chol(AR_mat))
    # Multiplying the transpose of the Cholesky factorization by a vector of random values gives vectors of autocorrelated value
    ar_var[[(((x-1-1)%/%12)+1)]] <<- L%*%matrix(rnorm(round(timescale)*(12^2)*round(timescale),0,1), ncol= round(timescale)*12)*r_SD*sqrt(1-phi^2)
  }
  random_values <- as.vector((ar_var[[(((x-1-1)%/%12)+1)]])[,1])[((((x-1-1)%%12)*round(timescale)+round(timescale)-round(timescale)+1)):(((x-1-1)%%12)*round(timescale)+round(timescale))]
  random_vec_summaries[[x-1]] <<- summary(exp(random_values - (sd(random_values)^2)/2))
  mapply ( function(growing) {
    # Total maximum energy consumptions across all fish of different sizes
    nu<-sum(size_dependent_energy_intake(structural_mass(new_E_l[[2]]), p_0, p_1)*new_mu_exp)
    # Compute a random value for the resource's carrying capacity
    random_value <- random_values[growing]
    random_value <- random_values[growing]
    random_resource_K <- resource_K*(exp(random_value - (sd(random_values)^2)/2))
    random_adjustment <- (exp(random_value - (sd(random_values)^2)/2))
    random_adjustment <- ((random_adjustment - 1)/1000)+1
    temperature <- Tref*random_adjustment
    p_0 <<- Arr(temperature, A_int)*p_0
    c_S <<- Arr(temperature, A_met)*c_S
    c_E <<- Arr(temperature, A_met)*c_E
    # Toggle which of these temperature dependencies are commented or not (turned off or on in the code) to have temperature have separate direct effects on each of these parameter values
    resource_K <<- Arr(temperature, A_car)*resource_K
    # resource_r <<- Arr(temperature, A_gro)*resource_r
    # m <<- Arr(temperature, A_mor)*m
    # Allow the fish to feed (with their maximum food intake dependent on their sizes the previous day) and the resource to simultaneously grow
    # Differential equation function
    Feeding <- function(t, state, parameters) {
      with(as.list(c(state, parameters)), {
        dR <- resource_r*R*((resource_K-R)/resource_K) - nu*((R)/(R+K_half))
        dC <- nu*((R)/(R+K_half))
        list(c(dR, dC))
      })
    }
    # Parameters for the DE
    parameters <- c(resource_r = resource_r, resource_K = resource_K, K_half=K_half, nu=nu)
    # States of resource density and resource consumed
    state <- c(R=resource, C=0)
    times <- seq(0, 1, by = 0.1)
    # Numerically integrate the differential equations
    out <- ode(y = state, times = times, func = Feeding, parms = parameters)  
    if (as.data.frame(out)$R[nrow(out)]<2) {
      print(out)
    }
    resource <<- as.data.frame(out)$R[nrow(out)]
    # Here, the fish grow
    new_E_l <<- get_pre_reproductive_size(new_E_l[[1]], new_E_l[[2]], resource, new_mu_exp, as.data.frame(out)$C[nrow(out)], nu)
    # New population sizes for each cohort-genotype combo, after mortality and prior to breeding
    before_fishing <- biomass_t(new_E_l[[1]], new_E_l[[2]], new_mu_exp)
    temp_new_mu_exp <<- fishing_mortality(new_mu_exp, new_E_l[[1]], new_E_l[[2]])
    after_fishing <- biomass_t(new_E_l[[1]], new_E_l[[2]], temp_new_mu_exp)
    yield <<- before_fishing - after_fishing
    new_mu_exp <<- mortality(new_mu_exp, new_E_l[[1]], new_E_l[[2]])
  }, 1:round(timescale))
  # Record resource level
  resources[x] <<- resource
  # Record consumption level
  # Resource consumption is only for last day within period of days in 'timescale' (i.e., the last day of the season if, timescale = 365/4)  
  resource_consumption[x] <<- new_E_l[[3]]
  # Record yield
  yields[x] <<- yield
  # Add oldest and second-oldest cohorts to make the 'plus group'	  
  new_mu_exp[nrow(new_mu_exp)-1,] <- new_mu_exp[nrow(new_mu_exp),] + new_mu_exp[nrow(new_mu_exp)-1,]
  # Empty out first row for young of year, for the mu_exp, E and l matrices
  new_mu_exp <- advance_age(new_mu_exp)
  new_E_l[[1]] <- advance_age(new_E_l[[1]])
  new_E_l[[2]] <- advance_age(new_E_l[[2]])
  # Empty out first row for young of year, for the mu matrix
  mu <- list[[4]]
  mu[2:nrow(mu),,] <- mu[1:(nrow(mu)-1),,]
  mu[1,,] <- 0
  # Reproduction is seasonal
  if ((breeding_season==T) & x%%round(365/timescale ) == 0 & x!=runtime) {
    # Get the amount of eggs produced and the amounts produced by each cohort-genotype combo, and a new E after reversible energy is depleted by mating
    fecundity_object <- fecundity(new_E_l[[1]], new_E_l[[2]], new_mu_exp)
    # The updated E matrix with cohorts' individuals' reversible mass values depleted (if they mated)
    new_E_l[[1]] <- fecundity_object[[3]]
    # Update first row of mu_exp with new individuals with expression based on past genotype
    new_recruits <- Ricker(fecundity_object[[1]])
    # Record the total number of new recruits
    recruitments[x] <<- new_recruits
    if (new_recruits>0) {
      inheritance_object <- inheritance(fecundity_object[[2]], mu, new_mu_exp, Ricker(fecundity_object[[1]]))
      # Updating mu_exp matrix
      new_mu_exp <- inheritance_object[[4]]
      # Give the YOY their sizes and lengths, if there are any
      if (sum(new_mu_exp[1,]) > 0) {
        new_E_l[[1]][1,] <- 0.3712037
        new_E_l[[2]][1,] <- 0.057
      }
      # Create vectors with length=number of new individuals, where each value is the individual's expressed phenotype (for "expressed") and their genotype (for "genotype")
      # "genotype" has one of these for each locus (for each locus, giving the alleles that each of the individuals have for that locus)
      mu <- inheritance_object[[1]]
      expressed <- inheritance_object[[2]]
      genotype <- inheritance_object[[3]]
      # length(expressed) should == length(genotype)
      # Filling in the genotypes of the new recruits
      # The scroll_through_genotypes function will create a row for each z-level in the new mu 3D array, where the row has the proportions of new recruits from each phenotype (column) that have the '1' allele for the locus that this z-level of the array corresponds to    
      # See "reproduction_function" file for more info in comments
      new_cohort_genotypes<-mapply(function(k) scroll_through_phenotypes(expressed, genotype[,k]), seq(1, loci))    
      mu[1,,] <- new_cohort_genotypes
    }
  }
  # Return updated matrices for this time point (2 through runtime)
  return((list(new_E_l[[1]], new_E_l[[2]], new_mu_exp, mu)))
}

# Function for advancing age
advance_age <- function(matrix) {
  matrix[2:(nrow(matrix)),] <- matrix[1:(nrow(matrix)-1),]
  matrix[1,] <- 0
  return(matrix)
}
```
