## Supplementary material for "Environmental variability alters and obscures fisheries-induced evolution": Code and simulated data files: mortality_functions.R

```
# MORTALITY FUNCTIONS

# Fishing mortality is size selective
selective_mortality <- function(length) {
  # q_f was renamed to z_f in the most recent manuscript draft
  1 / (1 + exp(-q_f*(length - mincatchsize)*100))
}

# Mortality drops as E/S increases, as S increases
mortality <- function(mu_exp, E, l) {
  lambda <- (E)/structural_mass(l)
  lambda[is.na(lambda)] <- 0
  mu_exp <- mu_exp*exp( -m - Fishing*selective_mortality(l) - m_c_max*exp(-z_c*lambda) - (m_p_max-m)*exp(-z_p*l))
  return(mu_exp)
}

fishing_mortality <- function(mu_exp, E, l) {
  lambda <- (E)/structural_mass(l)
  lambda[is.na(lambda)] <- 0
  mu_exp <- mu_exp*exp( - Fishing*selective_mortality(l))
  return(mu_exp)
}
```
