## Supplementary material for "Environmental variability alters and obscures fisheries-induced evolution": Code and simulated data files: readme.txt

This is the github repository for the paper "Food variation alters and obscures fisheries-induced evolution". 

Please see https://ecoevoenergymodels.shinyapps.io/Supplement/ for a description of the purpose of the code files in this repo. 

Please see the "smaller_files_figures_data_code" folder, and the "large_data_files" folder, in this repo for data (simulated values) used for figures and the code for plotting the figures. The files in "large_data_files" may be too large to be supported by the anonymized version of this repo. 

The four "model_run_code" files are for running the cod-like and herring-like versions of the model, and for each of these two options there is a file for the thermal simulations specifically, and a file which is just the base version of the model which directly varies fishes' food and does not simulate thermal scenarios. The other code files in the main repo directory hold necessary functions for running the model.
